# Structural and mechanistic studies on human LONP1 redefine the hand-over-hand translocation mechanism

**DOI:** 10.1101/2024.06.24.600538

**Authors:** Jeffrey T. Mindrebo, Gabriel C. Lander

## Abstract

AAA+ enzymes use energy from ATP hydrolysis to remodel diverse cellular targets. Structures of substrate-bound AAA+ complexes suggest that these enzymes employ a conserved hand-over-hand mechanism to thread substrates through their central pore. However, the fundamental aspects of the mechanisms governing motor function and substrate processing within specific AAA+ families remain unresolved. We used cryo-electron microscopy to structurally interrogate reaction intermediates from in vitro biochemical assays to inform the underlying regulatory mechanisms of the human mitochondrial AAA+ protease, LONP1. Our results demonstrate that substrate binding allosterically regulates proteolytic activity, and that LONP1 can adopt a configuration conducive to substrate translocation even when the ATPases are bound to ADP. These results challenge the conventional understanding of the hand-over-hand translocation mechanism, giving rise to an alternative model that aligns more closely with biochemical and biophysical data on related enzymes like ClpX, ClpA, the 26S proteasome, and Lon protease.

## Introduction

AAA+ (ATPases Associated with diverse cellular Activities) proteins are a large family of ATP-dependent enzymes that remodel diverse cellular targets via large-scale conformational rearrangements driven by energy derived from γ-phosphate bond hydrolysis.^1^ AAA+ proteins are essential in all domains of life, operating on proteins and nucleic acids to perform critical functions in diverse processes, such as cell cycle regulation, cargo transport, DNA transcription, translation, membrane fusion, and protein homeostasis.^2^ The AAA+ complexes that remodel proteins face unique challenges, given that proteins and protein assemblies display incredible diversity in composition, stability, conformations, and subcellular localization.^1-3^ Moreover, the AAA+ protein translocases must operate efficiently on an irregular and diverse polypeptide track due to the variability in size and chemistry of the canonical 20 amino acids and associated post-translational modifications. The majority of AAA+ protein remodelers form ring-like hexameric assemblies with non-enzymatic N-terminal domains that serve dual roles in substrate recruitment and regulating holoenzyme function.^1^ Over the past decade, high-resolution cryo-electron microscopy (cryo-EM) structures of AAA+ proteins have collectively suggested that these systems employ conserved mechanisms to bind and thread protein substrates through a central pore using ATP-powered conformational rearrangements.^1,3,4^ However, despite these advancements, the molecular details regarding motor function and the underlying regulatory mechanisms governing substrate processing within specific AAA+ subfamilies remain largely unresolved.

The Lon protease is a member of the HslU/ClpX/Lon/RuvB (HCLR) clade of AAA+ proteins, with a structure and functionality conserved from bacteria to eukaryotes.^5^ Lon assembles as a homohexamer, with each monomer containing an N-terminal substrate-binding domain (NTD), an HCLR clade ATPase domain, and a C-terminal protease domain (PD). The NTD can be further subdivided into a globular domain (NTD^GD^), a coiled-coil domain (CCD), and a 3-helical bundle domain (NTD^3H^) that is integrated into the AAA+ cassette (**Figure 1A**). Previous cryo-EM structures of bacterial and eukaryotic Lon proteases have highlighted a series of large-scale structural rearrangements associated with substrate engagement and degradation.^6-12^ In the absence of substrate, Lon adopts an open, left-handed lock-washer conformation (LONP1^OFF^), with all ATPase domains bound to ADP and the PDs in an inactive state. Upon binding of nucleotide and substrate, Lon transitions to a “closed” conformation (LONP1^ENZ^), with the ATPase domains adopting a right-handed helical arrangement with the conserved pore loops from ATP-bound subunits intercalating with the bound polypeptide in a residue-agnostic fashion (**Figure 1B, S1**). Transition to the closed form results in PD activation for processive degradation of substrates as they are translocated into the proteolytic chamber.^8^ These prior structural findings suggest that Lon proteases harness around-the-ring ATP hydrolysis to drive a hand-over-hand substrate translocation mechanism with a step size of two-residues per ATP consumed (**Figure S1**).^3^ The hand-over-hand model has gained popularity in the field as the translocation model not only for AAA+ proteases, but also broadly for many AAA+ enzymes.^13-20^ However, this mechanism remains debatable, as many of the biochemical and single molecule studies on AAA+ proteases are not consistent with a strictly sequential hand-over-hand mechanism.^21-23^

**Figure 1.**
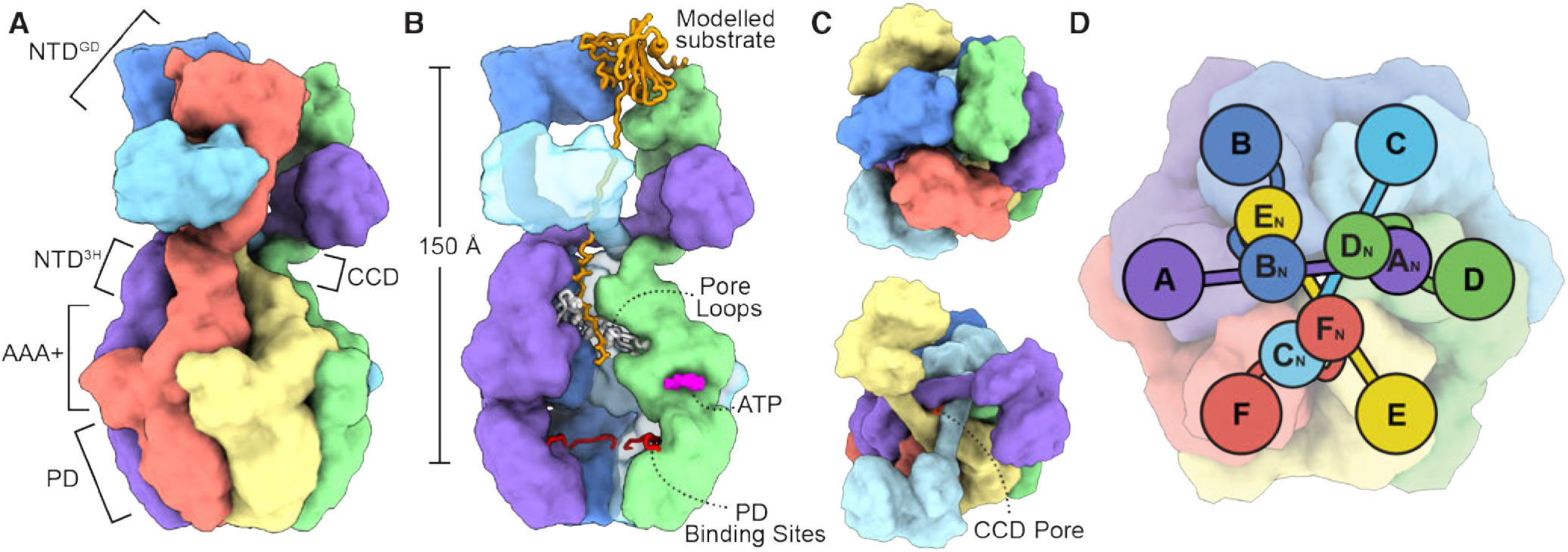
Overall architecture of LONP1 and NTD organization. **A**) Domain organization of LONP1 includes an N-terminal domain (NTD), a AAA+ ATPase domain, and a C-terminal protease domain (PD). The NTD is further subdivided into globular subdomain (NTD^GD^), a coiled-coil subdomain (CCD), and an NTD^3H^ subdomain that is integrated into the AAA+ cassette. **B**) Overview of structural elements required for substrate binding (modelled substrate) and translocation (white pore loops) as well as the nucleotide (magenta) and PD (red loops) binding sites. The distance allosteric signals must travel between the NTD to PD active site is provided on the left of the panel. **C**) The top panel shows a top view of the N-terminal domain trimer of dimer arrangement. The bottom panel shows a cutaway of the top three subunits to demonstrate the CCD pore positioned above the central axis. **D**) Cartoon representation of the overall organization of the NTD. The CCD pore is down the central axis of the 3-fold arrangement.

We recently structurally and biochemically investigated the specific activation mechanism of human LONP1,^6^ which plays a critical role in diverse mitochondrial processes such as proteostasis, metabolism, genome stability, and responses to hypoxic and oxidative stress.^24-29^ Our previous study showed that the left-handed open conformation of human LONP1 has a smaller pitch than bacterial homologs, and that human LONP1 is unable to fully adopt the proteolytically active closed-form in the presence ATPγS. Addition of the covalent PD inhibitor bortezomib was required to stabilize the active PD conformation (LONP1^BTZ^).^6^ Given that bortezomib is a peptide mimetic, this result led us to hypothesize that peptide substrates may play an important role in regulating holoenzyme activation. However, our inability to resolve the NTD in these prior reconstructions made it difficult to correlate substrate recognition with holoenzyme activation.

Substrate-engaged LONP1 complexes in multiple conformational states have also been determined by other groups,^7,30^ providing the first insights into the function and complex organization of the NTD, which organizes as a trimer of dimers via interweaving CCD helices (**Figure 1C**). The CCD of one protomer traverses the hexameric axis to dimerize with the CCD of the opposite protomer in the hexamer (e.g. position A to position D in **Figure 1D**), positioning the NTD^GD^ of one protomer above the other. Three repetitions of this pattern atop the hexamer positions long helices directly above the central axis to form a triangular pore (CCD pore), with the six NTD^GD^ adopting an alternating up-down arrangement (**Figure 1C**). This towering three-fold assembly is thought to be involved in coordinating substrate transfer from the NTD to the ATPase domains. This unique N-terminal structure is suggestive of allosteric communication between the NTD and the enzymatic domains of LONP1, whereby substrate recog-nition and unfolding at the NTD is coupled to the ATP-driven mechanochemical cycle. However, the molecular mechanism underlying such allostery has yet to be characterized.

Here, we used cryo-EM to determine the predominant structural conformations of LONP1 present during different stages of in vitro biochemical reactions to explore NTD-mediated allostery and motor function. We demonstrate that substrate binding to allosteric sites on the NTD regulates proteolytic activity independently of ATP, suggesting that substrate interactions play a critical role in long-range allosteric communication a distance of *∼*150 Å across the 600 kD holoenzyme assembly (**Figure 1B**). Surprisingly, our structural data shows that LONP1 can adopt a translocation-competent arrangement with the ATPases fully occupied by ADP. These data call into question key assumptions of the canonical hand-over-hand translocation mechanism, which requires strict coupling of nucleotide state to ATPase domain conformation. We also solved cryo-EM structures of LONP1 during and after in vitro proteolytic reactions to demonstrate that these structures represent stages of the dwell phase detected in numerous optical tweezer experiments.^31-33^ Our findings present an alternative to the canonical hand-over-hand mechanism that is more congruent with the biochemical and biophysical data on ClpX, ClpA, and Lon protease.^32-39^ This fresh perspective on the mechanisms governing AAA+ protease function provides insights into the complex regulatory systems governing LONP1 activity, and lays the groundwork for future studies investigating LONP1’s role in the regulation and maintenance of mitochondrial homeostasis.

## Results

### LONP1’s NTD is allosterically coupled to the PDs

Prior single particle cryo-EM studies defined the “trimer of dimers” arrangement of the Lon NTD, mediated by a complex interconnected network of helices derived from the CCD sub-domain (**Figure 1**).^7,9,10,30,40^ Although the NTDs have been implicated in regulating bacterial Lon protease activity,^41^ this relationship has not been demonstrated in human LONP1. We investigated NTD-mediated allostery with an established peptidase assay that uses a small fluorogenic peptide, Glutaryl-Ala-Ala-Phe-MNA (GAAF-MNA), as a readout for protease domain activity.^42^ Importantly, ATP-mediated translocation is not required to cleave this small peptidomimetic substrate. It was previously shown that LONP1 possesses a basal peptidase rate, and that addition of casein (a model substrate) leads to a two-fold peptidase rate enhancement.^43^ We reasoned that this assay could serve as a readout of substrate-induced PD activation. We evaluated LONP1 peptidase activity under four conditions: in the absence of substrates (basal rate), with casein, with ATP, and with both ATP and casein. In line with previous reports, the addition of casein alone led to a two-fold rate enhancement (**Figure 2A**). However, the addition of ATP had no effect on the peptidase rate, and the rate enhancement remained nearly two-fold in assays with both ATP and casein, suggesting that substrate, not nucleotide, allosterically activates the PD in the wt LONP1 system.

**Figure 2.**
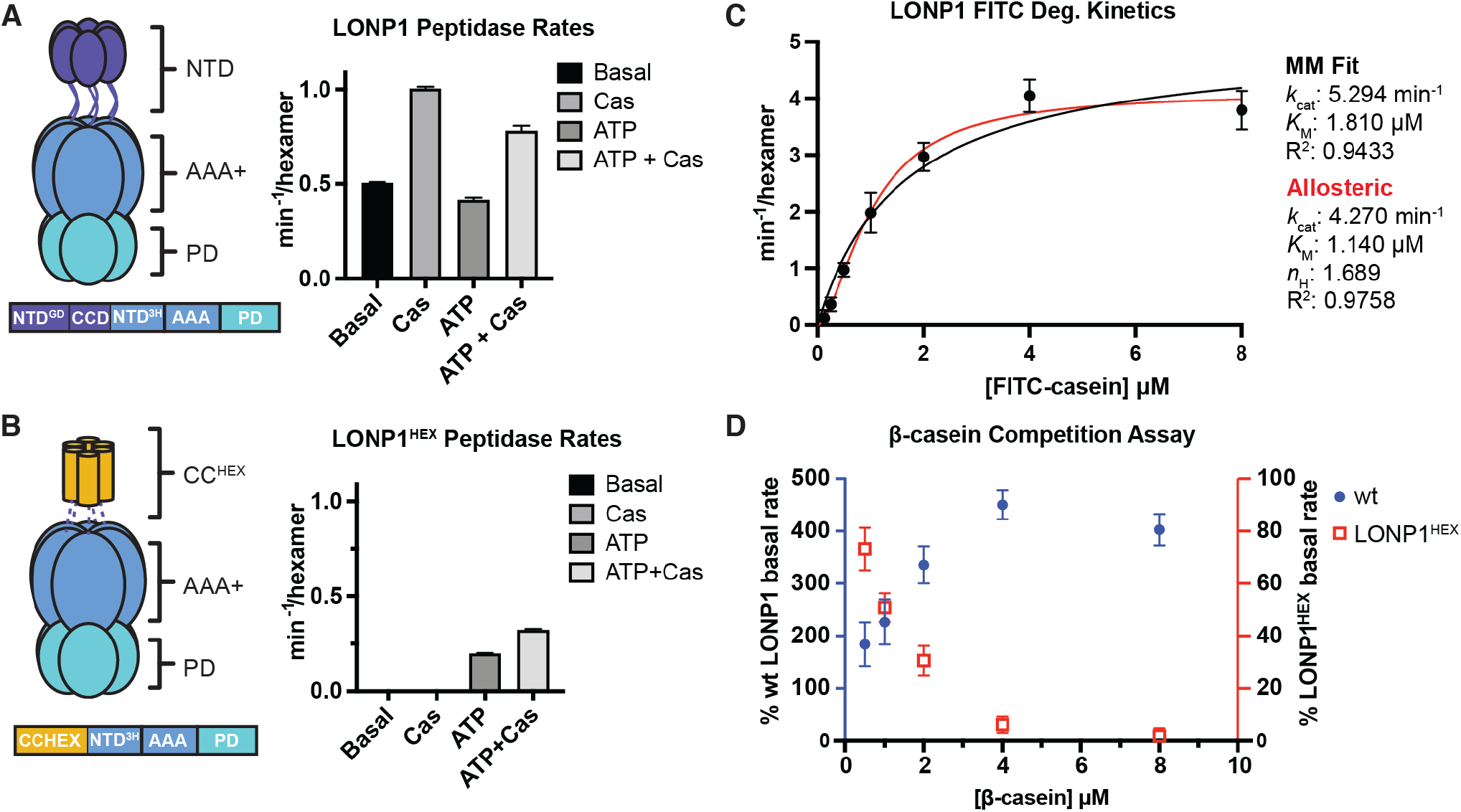
NTD-mediated allostery. **A**) wt LONP1 and **B**) LONP1^HEX^ peptidase assay results. The basal rate represents the rate of fluorogenic peptide cleavage without the addition of ATP or casein. **C**) FITC-casein Michaelis-Menten degradation kinetics shown with standard Michaelis-Menten non-linear regression (black line) or allosteric model (red line). Kinetic parameters for both models are provided on the right side of the graph. **D**) A coplot of LONP1 and LONP1^HEX^ β-casein competition assay results. In these experiments, FITC-casein was held constant at 1 µM and rates were determined in the presence of increasing concentration of unlabeled β-casein. All assays were repeated in biological triplicate and error bars represent the standard deviation on the mean.

We next aimed to more directly confirm the role of the NTD on ATPase and proteolytic activity. The NTD is crucial for LONP1 hexamerization,^40,44^ thus to evaluate the role of this region in allosteric activation of the PD we generated a LONP1^HEX^ construct where the NTD^GD^ and CCD were replaced by a previously established coiled-coil hexamerization domain (**Figure 2B**).^45,46^ The coiled-coil domain would preserve the hexamerizing influence of the LONP1 NTD, but would not contribute any substrate-recruiting capability. LONP1^HEX^ eluted from the size exclusion column (SEC) in a peak fraction corresponding to a hexamer, although its ability to hydrolyze ATP and degrade fluorescein isothiocyanate (FITC)-casein was significantly diminished, with rates 9-fold and 17-fold lower than wt, respectively (**Figure S2**).These data indicate that the NTD is intimately linked to LONP1 function, even in a construct where hexamerization of the complex is rectified with an additional synthetic domain shown to stabilize other AAA+ proteases for structure-function studies.^17,19,40,45,47^

With a functional NTD-truncated construct in hand, we next tested LONP1^HEX^ using our GAAF-MNA peptide assay. Surprisingly, LONP1^HEX^ did not have a detectable basal peptidase rate, and the addition of casein alone had no impact on activity (**Figure 2B**). However, the addition of ATP resulted in measurable peptidase activity, and the addition of both ATP and casein further augmented the rate (**Figure 2B**). This loss of substrate-stimulated peptidase rate enhancement in the absence of ATP in our LONP1^HEX^ construct suggests that substrate binding in LONP1’s NTD is allosterically coupled to the distal PD active sites. Further, the necessity of ATP to activate the LONP1^HEX^ complex suggests that substrate co-ordination with the ATPase domains facilitates the necessary conformational rearrangements to activate the protease in the absence of the NTD.

Given that our peptidase results are consistent with an allosteric governance of LONP1 function, we aimed to further support this model by measuring the kinetics of ATP-dependent degradation of FITC-casein. The allosteric model yielded a better fit than a Michaelis-Menten model, with a Hill coefficient of 1.69 (**Figure 1C**). Substrate-dependent allosteric activation of proteolytic activity was further supported by assaying LONP1 proteolytic activity against FITC-casein while titrating in increasing concentrations of unlabeled β-casein (**Figure 1D**). Under an allosteric model, low concentrations of the unlabeled substrate should increase degradation rates of the labelled substrate, whereas a non-allosteric model would be consistent with unlabeled substrate outcompeting the labelled substrate, resulting in decreased degradation rates with increasing concentration. We observe that unlabeled β-casein augments the rate of wt LONP1 in a dose-dependent manner, reaching a maximum rate enhancement 4.5-fold higher than the control at 4 µM β-casein. At this point, higher concentrations of casein begin to outcompete the labelled substrate, supporting an allosteric activation mechanism where substrate binding to an allosteric site leads to enhanced degradation rates. To verify that the allosteric binding sites are on the NTD, we then performed the same substrate competition assay using our LONP1^HEX^ construct. Unlike the wt enzyme, increasing unlabeled casein concentrations led to a steady decrease in LONP1^HEX^ activity (**Figure 1D**). Based on these results, we conclude that LONP1’s peptidase and proteolytic activities are allosterically coupled to substrate binding at the NTD.

### Purified LONP1 adopts a closed conformation in the absence of added substrate or ATP

Our peptidase assays indicate that substrate binding to LONP1 in the absence of nucleotide allosterically activates the distal PDs, suggesting that substrate interactions may be sufficient to drive a transition from the proteolytically inactive open conformation to the active closed conformation (**Figure S1**). To determine if the LONP1 used in our peptidase assay represents the inactive open-form of LONP1, we determined the cryo-EM structures of recombinantly expressed and purified wt LONP1 without adding nucleotide or proteolytic substrate. Surprisingly, over 90% of particles adopted the protease-active closed-form, with only a small minority of the particles adopting the open-form (**Figure S3**). Although the purified sample had not been supplemented with additional substrate, the closed-form reconstructions contain unmistakable substrate density traversing the center of the CCD pore into the AAA+ domain (**Figure 3A**). Similar substrate-engaged conformations have been identified in other studies, but only upon supplementation with nucleotide and substrate.^6,7,30^ Importantly, this observed substrate density bridging the NTD and the ATPase domains provides an additional structural tether between the NTD and ATPase domains that likely aids in facilitating allosteric communication across the assembly. This unexpected adoption of the closed-form conformer in the absence of added nucleotide explains LONP1’s basal peptidase rate as it is predominantly in the proteolytically active, closed conformation in our assay. Moreover, with LONP1 predominantly bound to substrate, these data demonstrate that the substrate-mediated rate enhancement of peptidase activity is due to casein associating with allosteric bindings sites on the NTD distinct from the main substrate binding channel.

**Figure 3.**
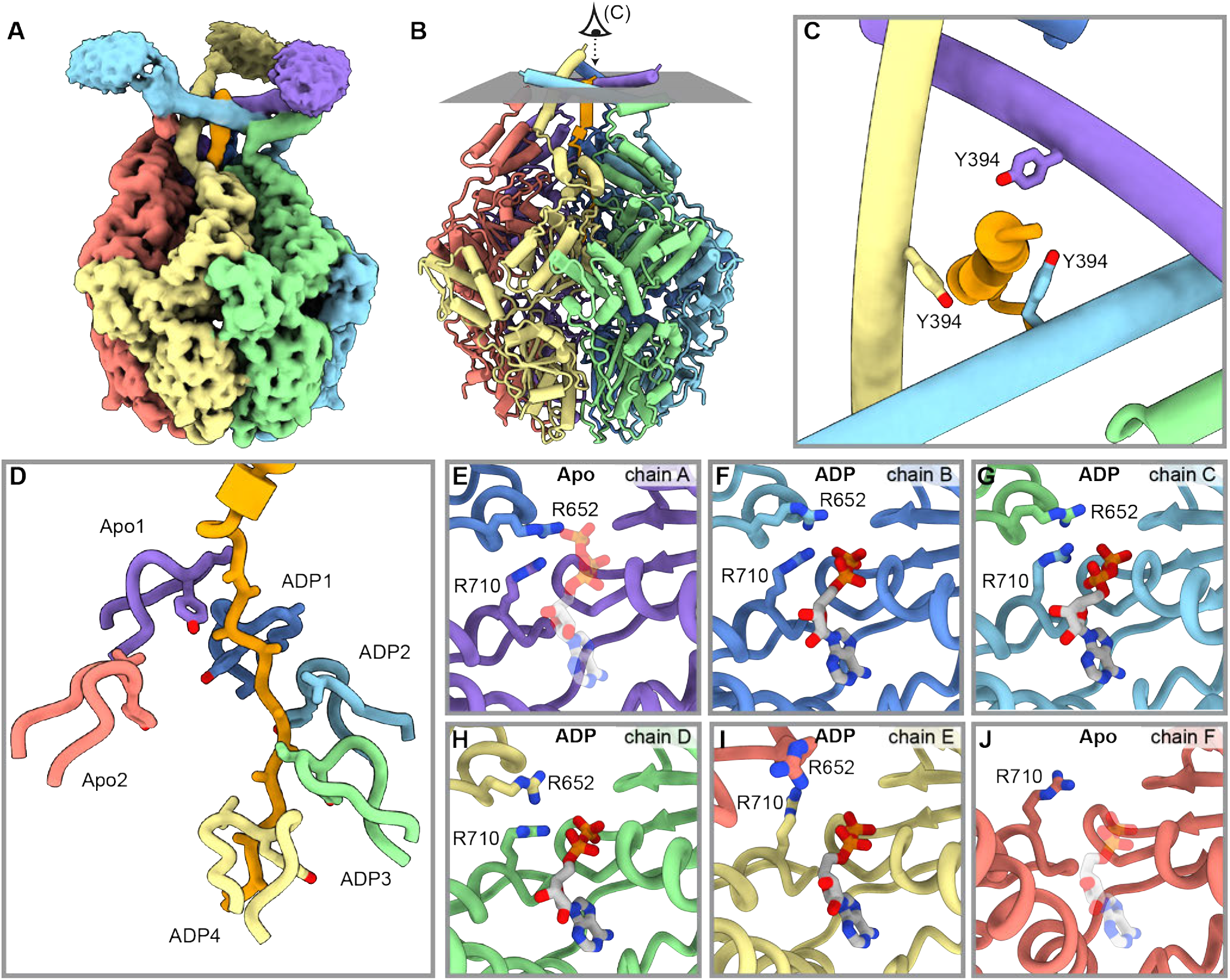
Structural overview of substrate-bound, closed form of LONP1 bound to ADP. **A**) Unsharpened cryo-EM map of substrate-bound LONP1 colored by chain demonstrating density for the CCD subdomain and bound substrate (orange). **B**) Atomic model of LONP1with viewing eye reference for panel (C). **C**) Substrate coordinated by Y394A from the CCD subdomain. **D**) Pore loops form a right-handed spiral around the proteolytic substrate and intercalate every other residue in the same manner as LONP1^ENZ^. The corresponding ATPase domain nucleotide state is labelled next to its respective pore loop. **E-J**) Nucleotide state of the ATPase active sites of positions A-F and conformational state of the cis- and trans-acting arginine fingers. Positions A and F are in an apo state (semi-transparent ATP for position A and semi-transparent ADP for position F are shown for reference to canonical active form, LONP1^ENZ^) while all other ATPase domains are bound to ADP.

Our peptidase assay and structural data show that substrate binding to the NTDs stimulates peptidase activity independent of nucleotide. To more thoroughly interrogate substrate-induced activation, we collected a cryo-EM dataset of LONP1 incubated with excess casein in the absence of additional nucleotides. The data again revealed predominantly the proteolytically active closed-form, but with improved interpretability of the CCD and peripheral NTD^GD^ (**Figures S4, S5**). Importantly, these results indicate that the addition of excess substrate in our sample leads to a more organized NTD arrangement, likely due to substrate association with allosteric binding sites. Using our improved maps, we built an atomic model of the assembly (**Figure 3B**). Although the CCD density was less pronounced due to the flexibility of this region, our best model positions the conserved Y394 at the center of the triangular CCD pore, extending toward the central axis to interact with the bound substrate (**Figure 3C, S6A**). The diameter of the substrate density at the center of the CCD is consistent with an α-helix, and the density becomes thinner and better resolved as it extends into the ATPase domains, where it is coordinated as an extended peptide by the conserved pore loops (**Figure 3D, S6B**). The position of the conserved Y394 at the center of the CCD pore and putative association with substrate is consistent with previous studies, suggesting that this residue is important for substrate processing in bacterial and metazoan Lon proteases.^9,30^

To further investigate the functional importance of Y394, we mutated this residue to alanine and performed NADH-coupled ATPase and FITC-casein degradation assays. The kinetic parameters for basal and stimulated ATPase activity are nearly identical for Y394A and wt LONP1, and FITC-casein degradation assays indicate that the k_*cat*_ for Y394A is roughly double that of wt LONP1 with an only marginally elevated K_*M*_ value (**Figure S7**). Y394 thus does not appear to be coupled to ATPase activity and only modestly alters proteolytic activity, yet its conservation from bacteria to eukaryotes, and its position at the center of the CCD pore, indicate that Y394 likely serves an important but nuanced role in substrate processing (see *Discussion*).

### Nucleotide state does not strictly determine AAA+ conformation in LONP1

Current mechanistic models of AAA+ protein translocases posit that the γ-phosphate of ATP is critical for coordinating the inter-domain interactions that stabilize a spiral staircase conformation that positions pore loops for substrate engagement. Numerous studies consistent with this model have led to the dogma that nucleotide state defines ATPase domain conformation.^3,17,48^ Consistent with this model, we previously observed that the inactive open-state, LONP1^OFF^, was fully ADP-bound, while the closed-state, LONP1^ENZ^, was ATP-bound.^6^ Moreover, only ATP-bound subunits strongly engaged with substrate, as seen in other AAA+ structural studies.^6,8^ We were thus surprised to observe that our casein-supplemented LONP1 structure, while nearly identical to prior structures of LONP1 in ATP-bound conformations, only contained ADP nucleotide (**Figures 3E-J, S8-9**). Notably, our ADP-bound closed form of LONP1 presents all the structural features previously ascribed to the ATP-bound state. The ATPases are arranged as a right-handed spiral staircase (chains A-E from uppermost to lowest) with a “seam” subunit (chain F) that is displaced from the spiral positioned at roughly the same height as chain A, the upper-most ATP bound protomer. Accordingly, the pore loop interactions with bound substrate show a similar pattern of engagement as previous structures, with the pore loops from the five spiraling ATPases intercalating into the centrally located substrate (**Figure 3D**).

The position of each ATPase within this conserved configuration, including pore loop interactions with substrate, is thought to be structurally linked to the presence or absence of a nucleotide γ-phosphate within the ATP binding pocket of each subunit. However, our closed-state ADP-bound LONP1 reconstruction upends this notion, as the ATP binding pockets of each subunit, including the positions of the trans-acting arginine fingers, are identical to previous ATP-associated closed-state structures **(Figure 3E-J**). The arginine fingers are thought to contact the γ-phosphate of the neighboring ATPase binding pocket and catalyze nucleotide hydrolysis, and for this reason their positions within the ATP binding pocket have been used by our group and others as a proxy for assigning the nucleotide state.^14,17,18^ Our LONP1 structure highlights that nucleotide state, or even the presence of nucleotide, does not solely dictate the ATPase architecture. Strikingly, the binding pocket of the uppermost ATPase of the spiral staircase (chain A) is consistent with previously determined ATP-bound subunits, yet is devoid of any nucleotide density **(Figures 3E, S9**). Chains B-D of the spiral contain unambiguous ADP density, yet the binding pocket arginine fingers are in canonical ATP-interacting positions. Only the binding pockets of chains E and F are consistent with previously determined structures of Lon, where chain E is bound to ADP with the trans-acting arginine fingers in a retracted position, and chain F without discernible nucleotide density (**Figures 3E-J, S8**). These data suggest that LONP1’s ATPase active site conformation is influenced by substrate engagement and the relative position in the helical register, rather than being strictly dictated by nucleotide state. Importantly, these data indicate that current models of the hand-over-hand mechanism over-emphasize the importance of nucleotide state, thus unnecessarily limiting the proposal of alternate or modified mechanistic models that can more accurately integrate the available structural and biochemical data.

Our observation that chain A adopts an ATP-bound conformation despite a complete absence of nucleotide density is particularly intriguing (**Figures 3E, S9**). Chain A appears poised to bind ATP at the top of the register to initiate subsequent rounds of nucleotide exchange, but the current hand-over-hand model posits that conformational changes are driven by ATP hydrolysis, which strictly occurs in the D position of the register. This raises an important question: if ADP is in the putative hydrolysis position, how is force generated to shift the helical register to enable continued nucleotide exchange after the uppermost subunit is occupied? Each register shift of the ATPase is associated with a two-residue translocation of the substrate, thus three additional register shifts, wherein nucleotide exchange occurs at the top position, would be required to transition to the traditional ATP-bound conformation. Such a transition would effectively translocate substrate six residues before an ATP-bound subunit is in the putative hydrolysis position to initiate hand-over-hand substrate translocation (**Figure 4**). These observations raise important questions regarding the widely accepted aspects of the canonical hand-over-hand mechanism and indicate that ATP binding and pore-loop interactions with substrate, in the absence of ATP hydrolysis, can facilitate register-shifting conformational rearrangements that contribute to the overall mechanochemical cycle.

**Figure 4.**
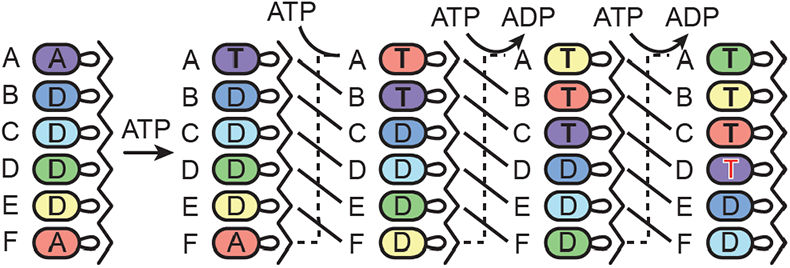
Proposed transitions resulting from nucleotide exchange. Linear representation of the hexameric spiral with D (ADP) and T (ATP) denoting the nucleotide state of each position, and A as empty. The putative hydrolysis position, position D, is shown with a red T when occupied with ATP.

### The ADP-bound closed structure represents the putative dwell state in the mechanochemical cycle

Our structural data show that recombinantly expressed LONP1 purifies with bound substrate, likely derived from an endogenous protein engaged during cell lysis and retained throughout the subsequent purification. LONP1 can access cellular ATP for such substrate processing during expression and cell lysis, but a lack of ATP in our purification buffers renders our ADP-bound closed-state reconstructions analogous to a stalled motor that has expended its fuel supply. This stalled state will hereafter be referred to as LONP1^stall^. The underlying properties of motor function of ClpXP, ClpAP, and Lon protease have been extensively characterized using optical tweezers, which collectively and consistently show that these motors alternate between dwell and burst phases representing 97% and 3% of the mechanochemical cycle, respectively.^32,34-37^ Nucleotide exchange occurs during the dwell phase while the burst phase consists of near-simultaneous ATP hydrolysis and/or phosphate release, resulting in force generation and rapid substrate translocation. Therefore, we reasoned that LONP1^stall^ could represent an early dwell-phase intermediate that occurs before subsequent rounds of nucleotide exchange. To test this hypothesis, we acquired cryo-EM data on LONP1 after incubation with ATP and casein for 30 minutes at 37 °C to ensure complete consumption of substrate and ATP. Our analyses identified both the open- and closed-state conformations of LONP1 (*∼* 33% and *∼* 67% of the particles, respectively) (**Figure S10**). The higher proportion of open-state particles compared to datasets without nucleotide supplementation confirms that the open conformation likely occurs after complete degradation of proteolytic substrate, as previously suggested.^6,8^

Further refinement of the closed-state resulted in a *∼* 2.9 Å reconstruction that was consistent with LONP1^BTZ^ (0.971 Å RMSD) and LONP1^stall^ (1.152 Å RMSD). However, we were surprised to observe ADP density present in all six of the active sites, including position A as well as the seam-subunit, position F (**Figure S11**). A comparison of the nucleotide density in position A with LONP1^ENZ^ indicates that, despite having nearly the same conformation and chemical environment, there is no discernible γ-phosphate density. Considering that LONP1^stall^ also adopts a similar active site arrangement despite lacking nucleotide density, this new all ADP-bound structure further supports the notion that nucleotide state does not determine ATPase active site conformation (**Figure 5**). These data suggest that this ADP-saturated closed state, which we refer to as LONP1^idle^, represents the initial conformation and nucleotide state of the putative dwell phase of the mechanochemical cycle before successive rounds nucleotide exchange. Furthermore, LONP1^stall^, which has nearly an identical conformation, thus represents this same dwell phase. We posit that during purification, substrate-engaged LONP1 retains four tightly bound ADP molecules while nucleotides in the seam and chain A binding pockets can dissociate. The lower nucleotide affinity for subunits at the top of the spiral confirms findings in previous studies that they are the site for nucleotide exchange,^3^ and that ATP binding necessitates register-shifting conformational rearrangements to position the next subunit for nucleotide exchange (**Figure 4**).

**Figure 5.**
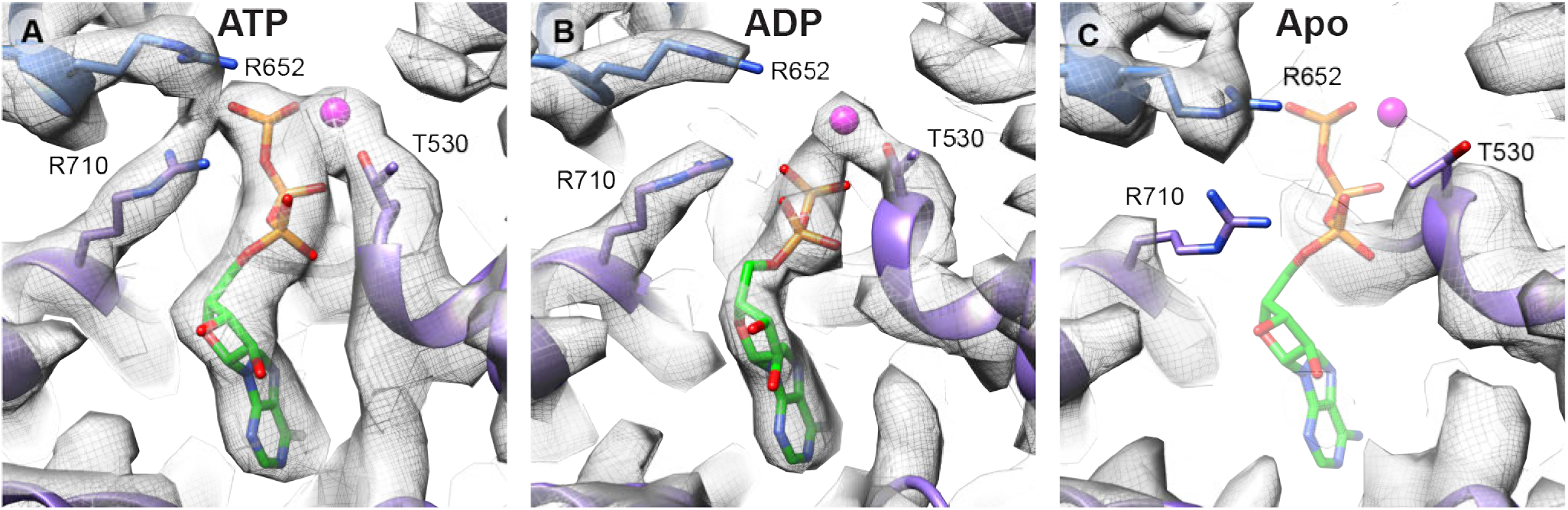
Comparison of nucleotide EM density in the A position for LONP1^ENZ^, LONP1^Idle^, LONP1^Stall^. **A**) ATP density in the A position for LONP1^ENZ^ represents the traditional ATP-bound state with strong γ-phosphate density coordinated to the two trans-acting arginine fingers. **B**) Nucleotide density in LONP1^Idle^ lacks density for the γ-phosphate but still retains the canonical conformation of the transacting arginine fingers in the presence of ADP. **C**) LONP1^stall^ lacks nucleotide density in the A position but still retains the same general conformation as LONP1^ENZ^ and LONP1^idle^. A semi-transparent ATP molecule is shown for reference.

### LONP1 represents an ensemble average of nucleotide states

Our structural data demonstrate that in LONP1, ATP does not strictly mediate ATPase domain conformations, in stark contrast to the current model of AAA+ protein translocases. Rather, ATPase conformation may be a register-specific phenomenon dictated by substrate engagement. Importantly, if nucleotide does not strictly define arrangement of the ATP binding pocket, then a structure of wt LONP1 actively degrading native substrates could allow the identification of ATPase states that do not fit the canonical hand-over-hand model. To test this hypothesis, we prepared cryo-EM grids of wt LONP1 degrading β-casein by rapidly plunge-freezing reactions after adding ATP. Data collection and subsequent processing readily identified the open and closed states. Further classification and refinement of the closed form resulted in a *∼* 3.3 Å reconstruction identical to our previous LONP1^BTZ^ structure (**Figures S12, S13**). All six PDs are catalytically active and have discernible density for substrate in the proteolytic active sites, indicating that this state represents the fully activated LONP1 holoenzyme (LONP1^ENZ^) (**Figures S13, S14**). Substrate density is more pronounced in chains A-C where the ATPase domains occupy the first three positions of the register and less pronounced in the putative hydrolysis position (chain D) and ADP-bound subunits (chains E and F) (**Figure S14**). Therefore, substrate binding and/or peptide bond cleavage may be coupled to changes in ATPase domain conformation.

We next examined the nucleotide density in each of the ATPase domains, and noted that while density for the γ-phosphate was well-defined in the A and B positions, nucleotide in positions C and D showed weaker γ-phosphate density (**Figure 6A-D, S15**). We thus directly evaluated nucleotide state by measuring the presence of γ-phosphate density in each of the four active sites proposed to be ATP-bound according to the hand-over-hand translocation model.^6,7^ Notably, we observe a gradient of γ-phosphate density across the four ATPase binding pockets, with positions A and B having the highest signal and position D having the lowest signal (**Figure 6E,F**). As a control, we performed the same analysis on the trans-acting arginine finger, R652, which was constant across chains A-C and only drops in position D. Moreover, the same pattern of normalized R652 density for chains A-D is observed in ADP-bound states, LONP1^stall^ and LONP1^idle^, further indicating that the γ-phosphate is not strictly required to position the trans-acting arginine finger. These data suggest that our structure of LONP1 actively degrading casein is an ensemble average of particles with varied ATP/ADP occupancy. Importantly, prior kinetic studies have shown that Lon protease has two high-affinity and two low-affinity ATP binding sites, which roughly match the γ-phosphate distribution observed in our EM maps.^49,50^ Given that the putative dwell phase represents 97% of the mechanochemical cycle, LONP1^ENZ^ likely reflects a later stage of the dwell phase, where variable numbers of ATP molecules are associated within the hexamer in preparation for the burst phase.

**Figure 6.**
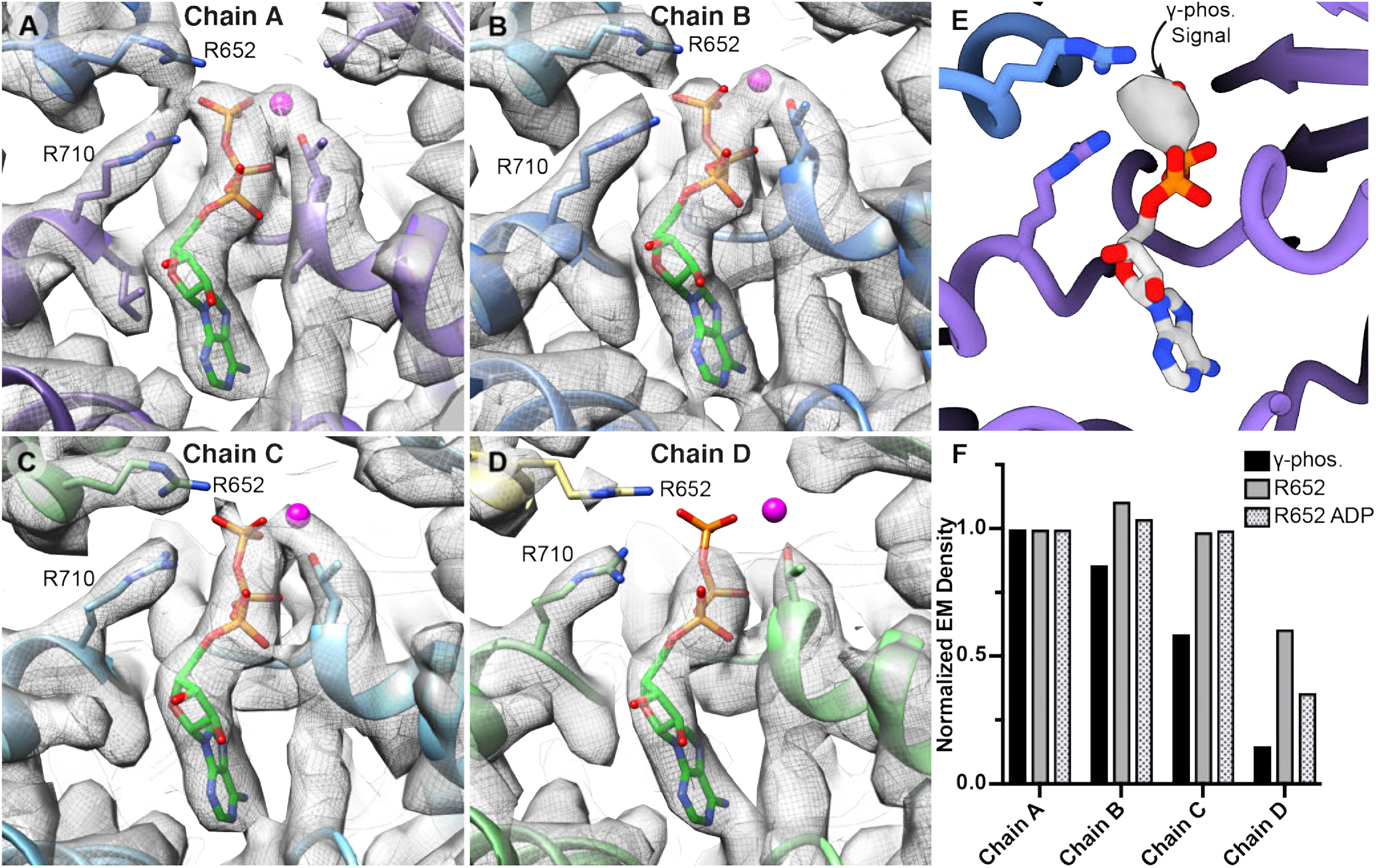
Actively translocating LONP1 represents an ensemble of nucleotide states. **A-D**) Nucleotide density in ATPase active sites of putative ATP-bound positions A-D. **E**) Example of a zoned map selection centered on the ATP γ-phosphate used to quantify the normalized ATP γ-phosphate densities in panel (F). **F**) Normalized γ-phosphate and trans-acting arginine finger (R652) density plotted by chain. All values are normalized to the density calculated from chain A. All calculations were performed at the same binarization threshold (0.35).

## Discussion

### NTD-mediated allostery in LONP1 function

A conserved feature of AAA+ proteases is the presence of an N-terminal domain that mediates substrate recruitment and processing directly or via adaptor proteins. While allosteric activators, and at least one substrate-specific adaptor protein, have been identified for bacterial Lon protease, to our knowledge there are no such reports for human LONP1.^51-55^ In the absence of adaptor proteins, the NTD must fulfill key regulatory functions regarding substrate recognition and processing. Recently published structures of LONP1 by two groups have defined the complex, interconnected assembly of the NTD that is suggestive of hardwired allosteric regulation in LONP1 (**Figure 1**).^7,30^ Despite these compelling observations, there are, to our knowledge, no biochemical studies directly demonstrating NTD-mediated allostery in human LONP1. To address this question, we employed biochemical assays to demonstrate LONP1’s distal proteolytic domains are activated by substrate binding to allosteric sites located on the NTD. Moreover, we further informed this allostery by structurally characterizing the conformational states present during our biochemical assays, which confirm the presence of allosteric interaction sites distinct from the main substrate binding channel.

An intriguing structural feature of our ADP-bound closed structures, which has also been observed in other Lon reconstructions, is substrate traversing the CCD pore and interacting with conserved tyrosine residue Y394 (**Figure 3C**).^7,9,30,34^ In this arrangement, substrate simultaneously contacts both the NTD and ATPase domains, coupling these distinct regions, assisting in allosterically regulating catalytic activity. However, the only difference observed for our Y394A CCD mutant was a modest 2-fold increase in K_*cat*_ for the degradation of FITC-casein while maintaining similar ATPase kinetics as the wt system. This raises a key question - What is the functional role of this residue if it seemingly decreases catalytic efficiency?

Recently, Kasal *et al*. showed that Mesoplasma florum Lon protease is not a particularly strong unfoldase, but it is the most processive thus far characterized.^36^ In their study, optical trap and biochemical studies confirmed that Lon protease degrades long multidomain substrates to completion without dissociating. Given that LONP1’s NTD is unique among AAA+ proteases with a CCD pore that contacts incoming substrate, we propose that this structural motif imparts Lon protease with its notable processivity. In support of this hypothesis, our structures confirm that wt LONP1 remains tightly bound to substrate even after extensive purification. Importantly, such conditions would likely lead to dissociation for other AAA+ proteases, which typically require the introduction of Walker B mutations or supplementation with ATP analogs to form stable complexes for structural studies.^3,48^

The LONP1 NTD was previously suggested to function as a molecular ratchet,^7^ which we can mechanistically detail with further complexity based our biochemical and structural data. We propose that the CCD pore contributes the pawl function necessary for the ratchet mechanism of the LONP1 ATPase, where around-the-ring hydrolysis is linked to a rotational motion of the CCD pore around the substrate to ensure unidirectional translocation during the translocation cycle (**Figure 7**). In this model, Y394 functions as a pawl to prevent backsliding and allow the pore-loops to reset and engage substrate with a minimized risk of dissociation. LONP1 would consequently continuously interact with the substrate at two distinct locations, thereby enhancing processivity by effectively increasing grip strength, a crucial characteristic of these motors.^56^ A two-component motor that links ATP-fueled power strokes to a ratchet and pawl would explain why Lon protease has a similar step size and translocation velocity as the double-ring ATPase ClpAP, rather than the larger step sizes and faster velocities seen for ClpXP.^32,57^ Moreover, removal of the pawl (Y394A) would likely decouple this dual-motor mechanism, enabling the ATPases to exert force unencumbered by the ratchet and pawl, leading to larger step sizes, faster degradation rates, and reduced processivity.

**Figure 7.**
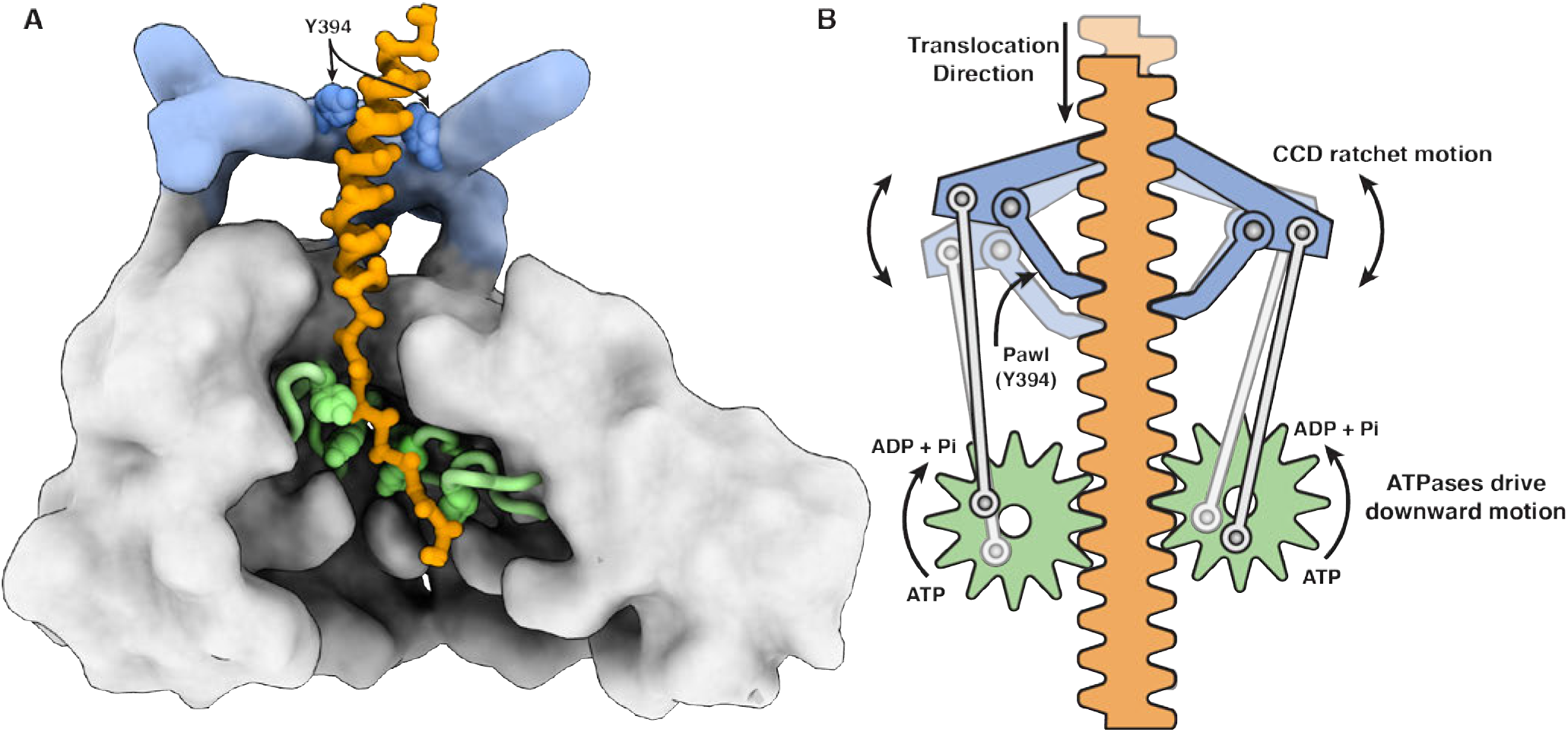
LONP1 ratchet and pawl translocation model. **A**) Cut-away of LONP1 engaged with proteolytic substrate. Interactions from both the CCD domain (light blue) of the NTD and the conserved ATPase pore loops (light green) engage substrate and participate in translocation. **B**) Cartoon model depicting LONP1 as a two-component motor combining a ratchet and pawl from the CCD with the hand-over-hand ATP fueled power strokes from the ATPase domains. The pawls (Y394) on the CCD facilitate unidirectional translocation as they ratchet around substrate due to conformational changes in the ATPase domains during hand-over-hand translocation.

### Relaxed hand-over-hand translocation model

Much of the available structural data for AAA+ proteases support a strictly sequential around-the-ring hand-over-hand mechanism (strict hand-over-hand), where each ATP hydrolysis event translocates two residues of a peptide substrate (**Figure 8A**).^3^ However, single-chain ClpX constructs can support substrate translocation for proteolytic degradation even with multiple catalytically dead ATPases.^39^ Such findings are at odds with a strictly sequential mechanism, as the motor would stall when the inactive mutant subunit reached the ATP hydrolysis position. Indeed, a growing number of biochemical and biophysical studies on ClpXP, ClpAP, and Lon protease are incongruent with a strict hand-over-hand interpretation.^32-39^ Optical tweezer experiments indicate that the fundamental step size for these motors is approximately six residues per ATP, and that kinetic bursts of up to 24 residues are possible.^31-33^ Kinetic bursts, even if limited to the fundamental six-residue step size, occur in rate regimes faster than the ensemble steady-state ATP hydrolysis rates of these systems, indicating fewer ATP molecules are hydrolyzed than required by a strict hand-over-hand model. Notably, the six-residue step size for Lon protease correlates with the time required to hydrolyze one ATP molecule, underscoring the likelihood of a six-residue step size per ATP consumed.^36^ Unsurprisingly, despite a broad acceptance of the hand-over-hand model, many aspects of the translocation mechanism of AAA+ proteins continue to be debated.

**Figure 8.**
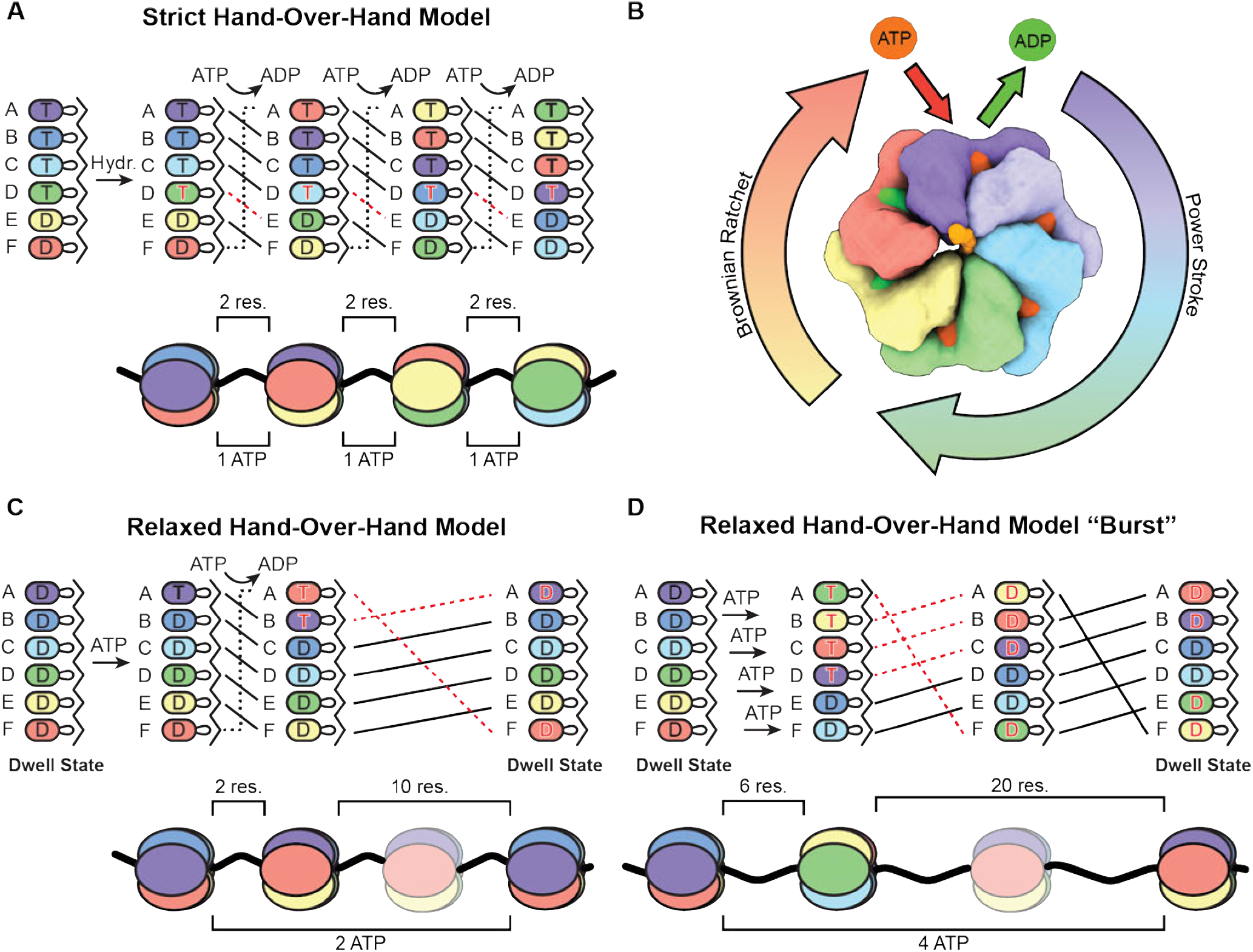
Proposed relaxed hand-over-hand translocation model. **A**) The strict interpretation of the hand-over-hand translocation mechanism where ATP hydrolysis occurs sequentially around the ring when subunits transition to the D position of the register resulting in a 2-residue translocation step size per ATP consumed (red dotted line indicates ATP hydrolysis). **B**) Overview of proposed energy-associated steps in the AAA+ translocation mechanism. Nucleotide exchange occurs at the top of the spiral (A position) and a power stroke leads to a compression of the angles between the large and small subdomains of the ATPase subunits. A return to the top of the spiral occurs via a Brownian ratchet derived from the relaxation of the compressed conformation of the large and small ATPase domains at the bottom of the spiral to a more extended conformation at the top of the spiral where pore loops reengage with substrate. **C**) Proposed relaxed hand-over-hand translocation mechanism that only undergoes nucleotide exchange at the top of the register when translocation stalls and the motor enters the dwell phase. Importantly, in this model, hydrolysis is not limited to the D position. **D**) Expansion of the relaxed translocation model to account for large bursts observed in optical tweezer experiments. In this interpretation, the ADP-bound AAA+ enzyme would continue coordinated hand-over-hand substrate translocation until kinetic energy is consumed and the motor stalls.

The strict hand-over-hand model is premised on nucleotide state defining conformation, which imposes limitations on mechanistic models that reconcile the available structural, biochemical, and biophysical data. However, our structural data demonstrate that a fully ADP-bound wt LONP1 can adopt a substrate-engaged conformation that is indistinguishable from ATP-bound LONP1^ENZ^, suggesting that pore-loop interactions with substrate may play a more salient architectural role than previously thought. Rather than passing through as a compliant cargo of the translocation mechanism, our data indicate that the polypeptide substrate plays a central role in dictating the hexameric arrangement. Thus, ATPase domain conformation appears to be a register-specific phenomenon.

Another limiting aspect of the strict hand-over-hand mechanism is the requirement for ATP hydrolysis to occur consecutively and exclusively at the lowest of the four ATP binding pockets that comprise the canonical ATPase staircase. Our data indicate that ATP binding at the topmost position of the staircase must shift the helical register regardless of the nucleotide state of the lowest ATP binding pocket, which is incongruent with the notion that ATP hydrolysis and nucleotide exchange are strictly coupled. Moreover, our ATP-bound LONP1^ENZ^ structure reflects an ensemble average of particles at different stages of the dwell phase, where the ATPase ring is filling with ATP. Under this interpretation, the partial occupancy of the γ-phosphate in the lowest binding pocket is more representative of an ensemble average of ATP and ADP bound states, as opposed to an ADP/Pi hydrolysis intermediate. Finally, the burst phase is likely a high-energy, conformationally dynamic state that rapidly resolves into the dwell phase and would represent only a small population of particles on the grid. From this perspective, the nucleotide state of LONP1^ENZ^ does not indicate that hydrolysis occurs exclusively in the lowest ATP binding pocket. However, the conformational transition of the ATPase to the lowest position is likely required for phosphate release after hydrolysis has occurred in subunits higher in the register, which would serve as the driving force for the power stroke.^32^

Our LONP1 data are inconsistent with a strict hand-over-hand mechanism for AAA+ protein translocation, leading us to propose a hybrid power-stroke and Brownian ratchet mechanism (**Figure 8B**).^58^ The power-stroke is coupled to ATP hydrolysis/phosphate release events that promote ATPase domains to successively progress through the registers of the spiral staircase from top to bottom. A power stroke, initiated by hydrolysis in one or more of the ATPase domains, coordinates the coupled, conformational transition of all the subunits within the ring, wherein each substrate-bound subunit participates in force generation regardless of nucleotide state, as demonstrated in single-chain ClpX studies.^35,59,60^ Transition to the bottom of the spiral compresses the angle between the large and small ATPase subdomains, storing some ATP-derived energy like a spring, held in position by the other five subunits. Once an ATPase subunit disengages from substrate at the bottom of the register, this stored energy drives the decompression of the subdomains to reposition the protomer at the topmost register of the ATPase staircase. Importantly, the ADP-bound seam subunit in LONP1^ENZ^ is found at a similar height as the ATP-bound subunit at the top of the spiral, demonstrating that this ratchet mechanism does not require ADP-dissociation.

Our structures indicate that nucleotide exchange occurs once an ATPase has assumed the topmost position in the spiral staircase, with its pore-loops re-engaged with the incoming substrate. However, our structures also show that dissociation of ADP, which is the rate limiting step of the dwell phase,^61^ is not required to enter the helical register and engage substrate during an active power stroke (**Figure 5**). Thus, it is conceivable that during a power stroke, an ADP-bound subunit positioned at the top of the staircase could progress through the spiral registers and participate in substrate translocation without undergoing nucleotide exchange, mediated entirely by association with the polypeptide substrate and neighboring subunits (**Figure 8C**). This efficient use of the kinetic energy generated by ATP hydrolysis avoids the requirement for time-consuming nucleotide exchange events at the top of the spiral. Nucleotide exchange would thus only occur during dwell phases of the translocation cycle, as observed in optical trapping experiments.^32,61^ We refer to this model as the relaxed hand-over-hand mechanism, as it still requires hand-over-hand substate translocation, but is uncoupled from the strict requirement for a step size of two residues per ATP hydrolysis. This model allows for “burst kinetics,” where coordinated ATP hydrolysis in one or more of the active sites fuels a two-residue hand-over-hand translocation mechanism without the need for continual nucleotide exchange or hydrolysis during each step (**Figure 8C**). This model, which is not strictly sequential nor stochastic, also accommodates the larger step sizes (e.g. six residues per ATP) measured by single-molecule studies. Previously reported translocation bursts of *∼*18-24 residues are also conceivable according to this model. Provided the kinetic energy generated from multiple coordinated ATP hydrolysis events is sufficient, and the linear sequence of amino acids provides adequate grip, the motor could feasibly continue sustained movement along the polypeptide track in an entirely ADP-bound state until molecular friction stalls the motor for the next dwell phase (**Figure 8D**).

Our relaxed hand-over-hand model is in line with a mechanism proposed for ClpX by Sen *et al*.,^32^ wherein nucleotide exchange occurs during fixed dwell phases and the total number of ATP nucleotides bound prior to the hydrolysis burst is determined by both ATP concentration and relative affinity of each nucleotide binding pocket for ATP. ClpX, like LONP1, has two high and two low-affinity ATP sites.^49,62^ In our actively translocating LONP1 dataset, we measured a distribution of ATP density that is consistent with this proposal, as the top two ATP binding pockets of the staircase have the strongest γ-phosphate density while the density decreases in the lowest two binding pockets (**Figure 6**). Furthermore, since the dwell phase represents 97% of the mechanochemical cycle, it is likely that all reported cryo-EM structures of AAA+ proteases bound to their respective substrates are snapshots of this phase, especially since the use of non-hydrolyzable ATP analogs or Walker B mutations likely impedes or prevents the burst phase.

While our relaxed hand-over-hand mechanism reconciles cryo-EM structures of AAA+ proteins with prior biochemical and biophysical studies, questions remain regarding the number of subunits that bind/hydrolyze ATP and the stochasticity underlying this process. Moreover, whether all ATP-bound subunits are required to fire during each cycle, or if these motors ever operate under a strict hand-over-hand translocation model, remain unknown despite these studies. It is tempting to suggest that many of the defining translocation parameters are dictated by the resistive force posed by the substrate, where more or fewer ATPase subunits bind and hydrolyze ATP depending on underlying factors such as local sequence identity or the presence of partially structured regions. Moreover, the unfolding of structured domains could represent an extreme case that necessitates the strict hand-over-hand mechanism, where maximal grip and controlled, successive hydrolysis encourages and rapidly traps destabilized thermal states. The field has not yet developed a capacity to precisely measure the kinetic parameters governing motor activity during the pre-unfolding dwell stage, requiring further innovations and collaborations to shed additional light on these important aspects of substrate processing. We thus hope that the structural data and model proposed herein provide new avenues to investigate the mechanisms driving these elegant molecular machines.

## Materials and Methods

### Construct generation

LONP1^CCHEX^ was designed in silico by removing the NTD sequence (deletion of residues 1-409) until the glycine linker (G410) that connects the CCD domain with NTD^3H^. Similar to previous studies,^45^ this region was replaced using a synthetic 32 residue hexamerization domain, connected by a flexible linker (GSGSYFQSNA) to the N-terminus of LONP1. The full construct was then synthesized and cloned into pET28a expression vector using NCO1 and XHO1 cut sites by Twist Biosciences. The Y394A LONP1 mutation was generated by site directed mutagenesis using the method developed Liu and Naismith.^63^ The resulting mutation was verified by sanger sequencing (Genewiz).

### Protein expression

LONP1 was transformed into Rosetta (DE3) PlysS cells (Sigma). 20 ml cultures were grown overnight and used to inoculate 1 L of terrific broth media (VWR). Cells were grown until they reached an OD of 0.8 and then cooled on ice before transferring to a 16 °C incubator for overnight induction using 0.5 M IPTG (Sigma). Cells were harvested by centrifugation and frozen until purification. To purify LONP1, cell pellets were resuspended in lysis buffer (50 mM Tris pH 8.0, 300 mM NaCl, 10% glycerol) to a volume of 40 ml. Cells were lysed by sonication and lysate was clarified at 30,000 *×* G for 45 minutes. Lysate was batch bound to nickel resin (Qiagen) for 30 minutes before applying to a column. The nickel beads were washed with 50 ml of lysis buffer, 50 ml of lysis buffer supplemented with 20 mM imidazole, and 50 ml of lysis buffer supplemented with 50 mM imidazole. Proteins were eluted off the column in lysis buffer supplemented with 250 mM imidazole. Fractions containing protein were pooled, concentrated, and injected onto a superose 6 increase 10/300 GL FPLC column (Cytiva) equilibrated with protein storage buffer (50 mM Tris pH 8.0, 150 mM NaCl, 0.5 mM TCEP). Fractions containing hexameric protein were pooled, concentrated, and flash frozen for future use.

### Peptidase Assay

Peptidase assays were performed by monitoring the cleavage of a commercially available fluorogenic peptide, Glutaryl-Ala-Ala-Phe-4-methoxy-β-naphthylamide (Sigma). For the basal peptidase rate, 1 mM of fluorogenic peptide was incubated in reaction buffer (100 mM KCl, 50 mM Tris pH 8.0, 1mM DTT, and 10 mM MgCl_2_) at 37 °C for 10 minutes before the addition of either LONP1 or LONP1^HEX^ at a final concentration of 100 nM. For reactions testing the effect of substrate on peptidase rate, the additional substrates (ATP, β-casein, or ATP + β-casein) were added to the reaction mixture and incubated with fluorogenic peptide for 10 minutes before the addition of enzyme. All reactions were monitored by an increase in fluorescence (excitation 340 nm, emission 430 nm) resulting from the liberation of 4-methoxy-β-naphthylamide using a Tecan plate reader. Data were fit using a linear regression and the slopes were used to determine raw rates. A standard curve of 4-methoxy-β-naphthylamide was used to convert fluorescence units into molar quantities. GraphPad Prism software was used for all data analysis. All reactions were repeated in triplicate and error bars from reactions represent standard deviation.

### Coupled ATPase assay

ATPase assays were performed by coupling the hydrolysis of ATP to the reduction of NADH by lactate dehydrogenase (LDH). Briefly, pyruvate kinase (PK) regenerates ATP from ADP produced during the reaction during the conversion of phosphoenol pyruvate (PEP) into pruvate. Pyruvate is then subsequently reduced in an NADH-dependent manner by LDH to produce lactate. The oxidation of NADH to NAD^+^ by LDH was monitored by a decrease in fluorescence (excitation 340 nm, emission 465 nm). Reactions were performed by first making a 2X mastermix of reaction components (2.0 mM PEP (1PlusChem), 0.4 mM NADH (Caymen Chemical), 60 U/ml LDH (Worthington Chemical), 20 U/ml PK (Sigma)). For activity measurements and kinetics, the mastermix was diluted to 1X in reaction buffer (100 mM KCl, 50 mM Tris pH 8.0, and 10 mM MgCl_2_) and incubated at 37 °C for 10 minutes with the appropriate concentration of ATP. Reactions were initiated by the addition of enzyme to a final concentration of 200 nM for wt LONP1 and Y394A or 500 nM for LONP1^HEX^). Data were fit using a linear regression and the slopes were used to determine raw rates. A standard curve of NADH was used to convert fluorescence units into molar quantities. Background NADH oxidation rates were determined by monitoring the reduction in fluorescence in the absence of enzyme and subtracted from rates determined in enzyme assays. Kinetic parameters were determined by fitting data using non-linear regression (Michaelis-Menten) in the GraphPad Prism software. All reactions were repeated in biological triplicate and error bars from reactions represent standard deviation.

### FITC-casein degradation assay

#### Activity Assay

FITC-casein (Sigma) was incubated with 2.5 mM ATP in reaction buffer (100 mM KCl, 50 mM Tris pH 8.0, and 10 mM MgCl_2_) at 37 °C for 10 minutes before initiating reactions by the addition of enzyme (250 nM LONP1, 2 µM LONP1^HEX^). An increase of fluorescence (excitation 485 nm, emission 535 nm) resulting from the liberation of FITC molecules was monitored using a TECAN plate reader. Three biological replicates were performed for each construct and the data were fit using a linear regression from which the slope was used to determine rate of substrate degradation. Data were normalized to wt LONP1 activity for comparison. All reactions were repeated in biological triplicate and error bars from reactions represent standard deviation.

#### β-casein competition assays

Assays were performed as described above, but in the presence of increasing concentrations of unlabeled β-casein (0.0 µM, 0.5 µM, 1.0 µM, 2.0 µM, 4.0 µM, 8.0 µM). All data were normalized to the basal rate in the absence of unlabeled substrate. All reactions were repeated in biological triplicate and error bars from reactions represent standard error of the mean.

#### Kinetics

Kinetics were determined as above but with varying concentrations of FITC-casein (0.125 µM, 0.250 µM, 0.500 µM, 1.000 µM, 2.000 µM, 4.000 µM, and 8.000 µM). Due to issues with the inner filter effect at higher substrate concentrations, FITC-casein was mixed in a 1:10 ratio with unlabeled β-casein to ensure accurate detection of liberated substrate. The resulting rates were corrected for the 10-fold difference before subsequent processing. The assays were initiated by the addition of 250 nM enzyme and fluorescence increase (excitation 435, emission 510) was monitored on a Tecan plate reader. A FITC-casein standard curve was generated by plotting the difference of initial and final fluorescence values determined after the complete degradation of known quantities of FITC-casein using bovine trypsin (Sigma). The standard curve was then used to convert fluorescence units into molar quantities. Kinetic parameters were determined by fitting data using a non-linear regression (Michaelis-Menten or allosteric model) in the GraphPad Prism9 software. All reactions were repeated in biological triplicate and error bars from reactions represent standard error of the mean.

### Sample preparation for Cryo-EM

For all samples, wild-type human LONP1 was diluted to appropriate concentrations in 50 mM Tris pH 8, 75 mM KCl, 10 mM MgCl2, 1 mM TCEP. Samples were incubated at 37 °C prior to vitrification using a Vitrobot Mark IV system (blot time 5 seconds, blot force 1, 100% humidity, room temperature, Whatman No. 1 filter paper). Four microliters of the sample were applied onto 300 mesh R1.2/1.3 UltrAuFoil Holey Gold Films (Quantifoil) that were glow discharged for 25 seconds at 15 mA with a Pelco Easiglow 91000 (Ted Pella, Inc.) in ambient vacuum. LONP1^stall^ samples were made by diluting LONP1 to 30 µM with or without 5 µM β-casein (Sigma) before applying to grids. For LONP1^idle^, 60 µM LONP1 was incubated with 30 µM β-casein (1:3 ratio LONP1 hexamer to substrate) and 2 mM ATP for 30 minutes at 37 °C. For LONP1^ENZ^, 30 µM of LONP1 was incubated with 5 µM β-casein (Sigma) for 10 minutes at 37 °C. Reactions were initiated by the addition of 2 mM ATP and immediately applied to grids and plunge frozen in 30 seconds.

### Cryo-EM data acquisition

Data for LONP1^stall^ (supplemented with casein) and LONP1^idle^ were collected on a Thermo Fisher Titan Krios operating at 300 keV and equipped with a Gatan K3 direct detection camera and a BioQuantum energy filter with a slit width of 20 eV. One-second exposures using an exposure rate of 30 e^-^/pixel/sec divided into 50 frames were acquired in counting mode, resulting in a total electron exposure of 54 e^-^/Å ^2^. The Leginon data collection software^64,65^ was used to collect micrographs at 105,000x nominal magnification (0.81 Å /pixel at the specimen level) with a set defocus of 1.0 µm. Stage movement was used to target the center of 49 holes and coma-compensated image-beam-shift^66^ was used to acquire high magnification images in the center of each of the 49 holes. A total of 4,580 micrographs were collected for LONP1^Closed^ and a total of 3,808 micrographs were collected for LONP1^Dwell^. Micrograph frames were aligned using MotionCor2 in the Appion image processing environment^64^ and processed in real-time using cryoSPARC Live^67^ to monitor image quality during data acquisition. See ***Table S1*** for details.

Data for LONP1^ENZ^ were collected on a Thermo-Fisher Talos Arctica transmission electron microscope operating at 200 keV equipped with a Gatan K2 Summit direct electron detector after setting parallel illumination conditions.^68^ *∼* 10 second exposures using an exposure rate of *∼* 7 e^-^/pixel/sec divided into ~50 frames were acquired in counting mode, resulting in a total electron exposure of *∼*50 e-/Å ^2^. The Leginon data collection software^64,65^ was used to collect micrographs at 36,000x nominal magnification (1.15 Å /pixel at the specimen level) with a defocus range of 0.9 µm to 1.6 µm. Stage movement was used to target the center of 16 holes and coma-compensated image-beam-shift^66^ was used to acquire high magnification images in the center of each of the 16 holes. A total of 1964 micrographs were collected. Micrograph frames were aligned using MotionCor2 and CTF parameters were estimated with CTFFind4 in real-time to monitor image quality during data acquisition using the Appion image processing environment.^64^ See ***Table S1*** for details.

### Image processing for LONP1^stall^, *LONP1*^idle^, *and LONP1*^ENZ^

The general processing workflow used for all datasets are described here, and further details are provided in the processing workflows in ***Figures S3-5, S9 and S11***. Movie frames were motion-corrected and dose-weighted using MotionCor2,^69^ and the summed and dose weighted micrographs were imported into cryoSPARC v2.1-v3.3.2^67^ for patch CTF correction. Micrographs with defocus values greater than 3.3 µm or with estimated resolutions worse than 7 Å resolution were discarded. Initial rounds of picking were done using a blob picker with an upper radius of 225 and lower radius of 125. For LONP1^idle^ and LONP1^stall^, particles were extracted using a 400-pixel size box and Fourier cropped to 256 pixels, while LONP1^ENZ^ was extracted in a 288-pixel size box without Fourier cropping. 2D classification was used to generate 6 templates (3 open form LONP1 and 3 closed form LONP1) that were then used for template picking with default settings and a particle diameter of 175 Å. Template-picked particles were extracted and lightly cleaned up using 2D classification. Particle stacks were then used in ab initio classification with default settings and three classes to generate initial open and closed models of LONP1. These were then used in a subsequent heterogeneous refinement job to separate out the open and closed conformations. Two closed form, 1 open form, and two “trash” classes were used during heterogenous refinement to identify false picks or damaged particles. The closed-form particle classes were then used for 3D classification using PCA initialization with default parameters except an O-EM learning rate initialization of 0.2. Classes were then refined using NU-refinement^70^refinement and the resulting models were evaluated and selecting using their resulting resolution, determined by an FSC of 0.143, and by visual interpretation of the refined maps. Final refinements were performed using NU-refinement^70^ with per particle defocus and CTF refinements enabled with standard settings. Reported resolutions were determined using the 3DFSC.^71^ Further details for the processing of each dataset can be found in the supplementary information.

### Atomic model building and refinement

All models of the closed-form LONP1 presented in this work were generated by first manually docking the previously determined LONP1^BTZ^ model (PDB:7KRZ)^6^ into the cryo-EM reconstructions using ChimeraX.^72^ Iterative rounds of model building and refinement were performed in PHENIX v1.19.2^73^ and Coot 0.9.4.1EL^74^ until reasonable agreement between the model and data were achieved. Final model relaxation and removal of clashes and bad rotamer outliers were performed using ISOLDE.^75^ Chimera^76^ and ChimeraX^72^ were used to interpret EM maps and models, as well as to generate figures.

### Calculation of γ-phosphate density

Using ChimeraX,^72^ we created zoned map selections centered on the ATP γ-phosphate from active sites A-D. We then integrated the total area from these selections and normalized them to the area from position A, which had the highest γ-phosphate density.

## Acknowledgements

We thank Will Lessin at the Scripps Research Electron Microscopy Facility for microscopy support. We thank Charles Bowman and J.C. Ducom at Scripps Research High Performance Computing core for computational support. This work is supported by the National Institutes of Health (NIH) NS095892 to G.C.L. and NIH F32GM145143 to J.T.M. Data collection used equipment supported by NIH grant S10OD032467. Computational analyses of EM data were performed using shared instrumentation funded by NIH S10OD021634.

## Author Contributions

JTM was responsible for conceptualization, methodology, data collection, investigation, formal analysis, validation, visualization, writing, funding acquisition (NIH F32) and much more than can be listed here. GCL helped out where he could, serving primarily as a cheerleader.

## Data availability

Cryo-EM maps and associated atomic models were deposited to the Electron Microscopy Databank (EMDB) and the Protein Databank (PDB), respectively, with the following EMDB and PDB IDs: LONP1l^ENZ^ - EMD-45430, 9BH6; LONP1^idle^ - EMD-45431, 9CC1; LONP1^stall+casein^ - EMD-45433, 9CC3; LONP1^stall^ - EMD-45434 (EMDB only).

## Supplementary Figures

**Figure S1.**
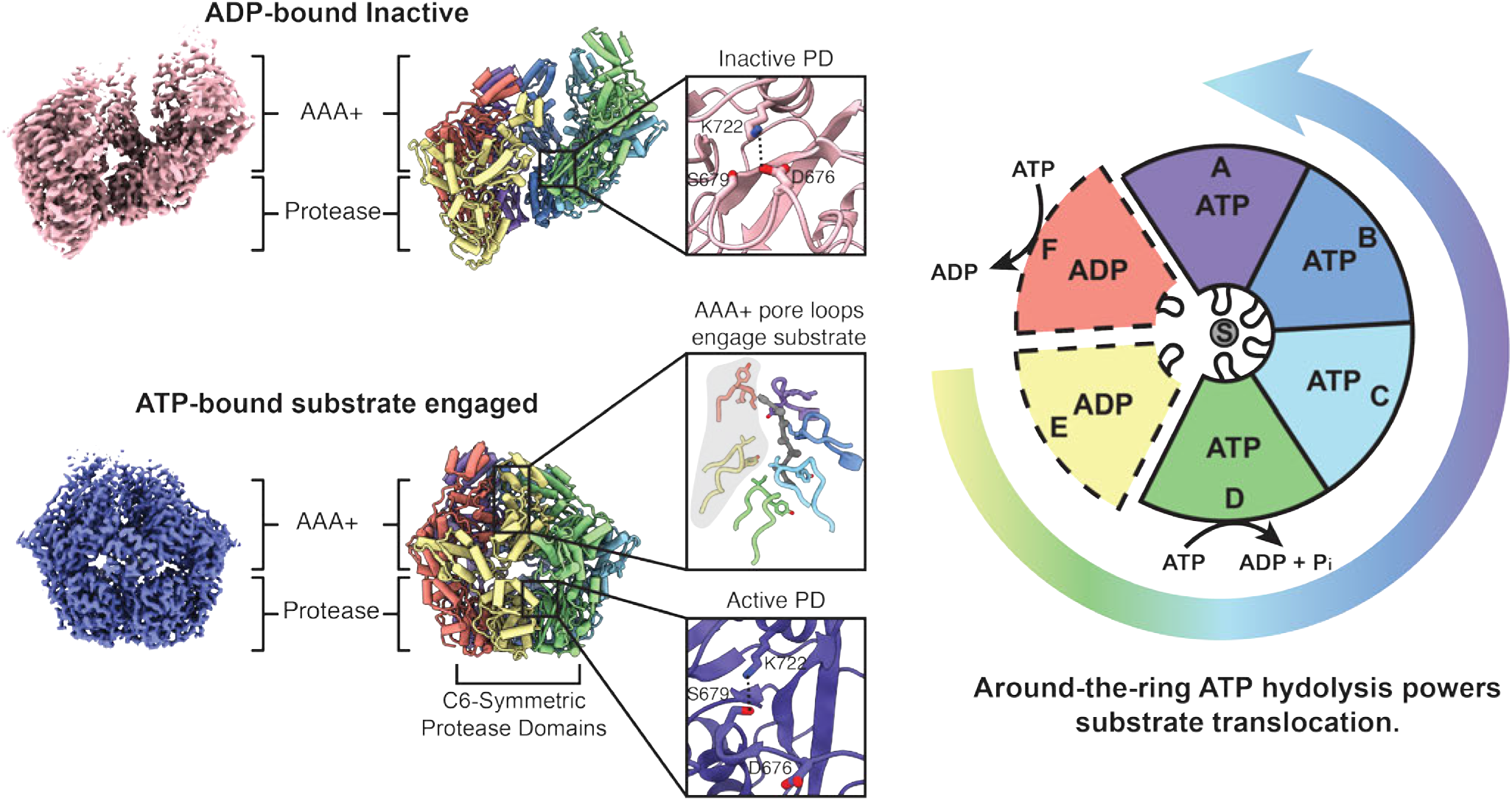
Activation of Lon protease and the hand-over-hand substrate translocation model. In the resting state, Lon protease is in a proteolytically inactive, open lockwasher conformation that forms a left-handed spiral (Top Left). In the open form (LONP1^OFF^), a 3_*10*_ helix is formed that positions a conserved aspartate adjacent to the catalytic lysine, blocking substrate access and neutralizing the catalytic base. Upon binding substrate and nucleotide, Lon transitions to the closed form where the conserved pore loops of ATP-bound ATPase subunits intercalate with the bound polypeptide every other residue (Bottom Left). Transition to the closed form results in a C6-symmeteric conformation of the CTDs where the inhibitory 3_*10*_ helix adopts a linear conformation that forms the substrate binding pocket. Lon proteases are proposed to utilize a hand-over-hand substrate translocation mechanism that couples around-the-ring ATP hydrolysis with conformational rearrangements to translocate substrates with a two-residue step size per ATP consumed (Right).

**Figure S2.**
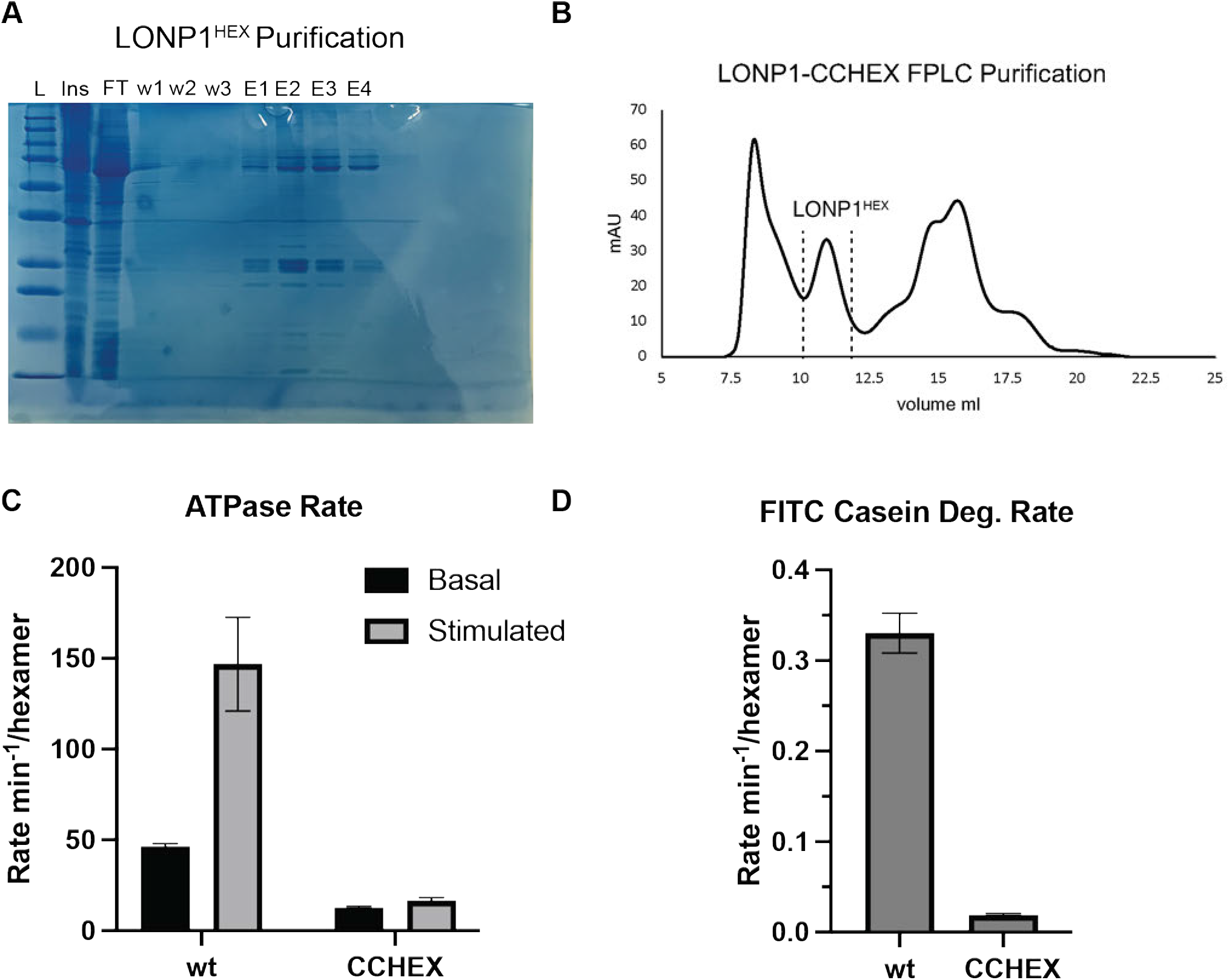
Purification and characterization of LONP1^HEX^. **A**) SDS-PAGE gel of LONP1 purification with the insoluble fraction (Ins), flowthrough (FT), washes fractions W1-W3, and elution fractions E1-E4. **B**) FPLC trace of pooled E1-E4 fractions run on a superose 6 increase 10/300 gl column. **C**) Basal and stimulated ATPase rates of wt LONP1 and LONP1^HEX^. Assays were run in triplicate and error bars represent standard deviation on the mean. **D**) FITC-casein degradation assay results of wt LONP1 and LONP1^HEX^. Assays were run in triplicate and error bares represent standard deviation of the mean.

**Figure S3.**
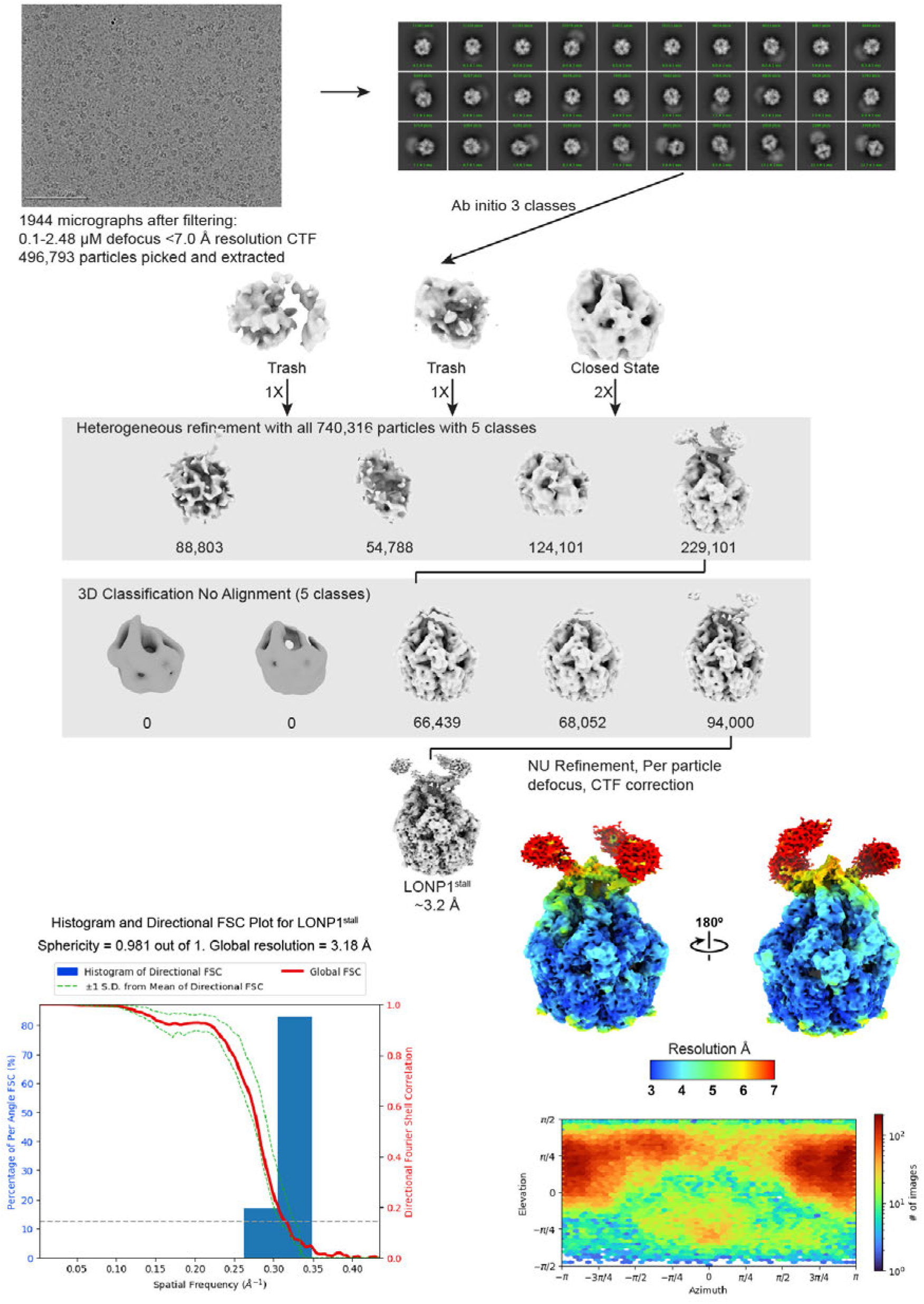
LONP1^idle^ no added substrates processing workflow.

**Figure S4.**
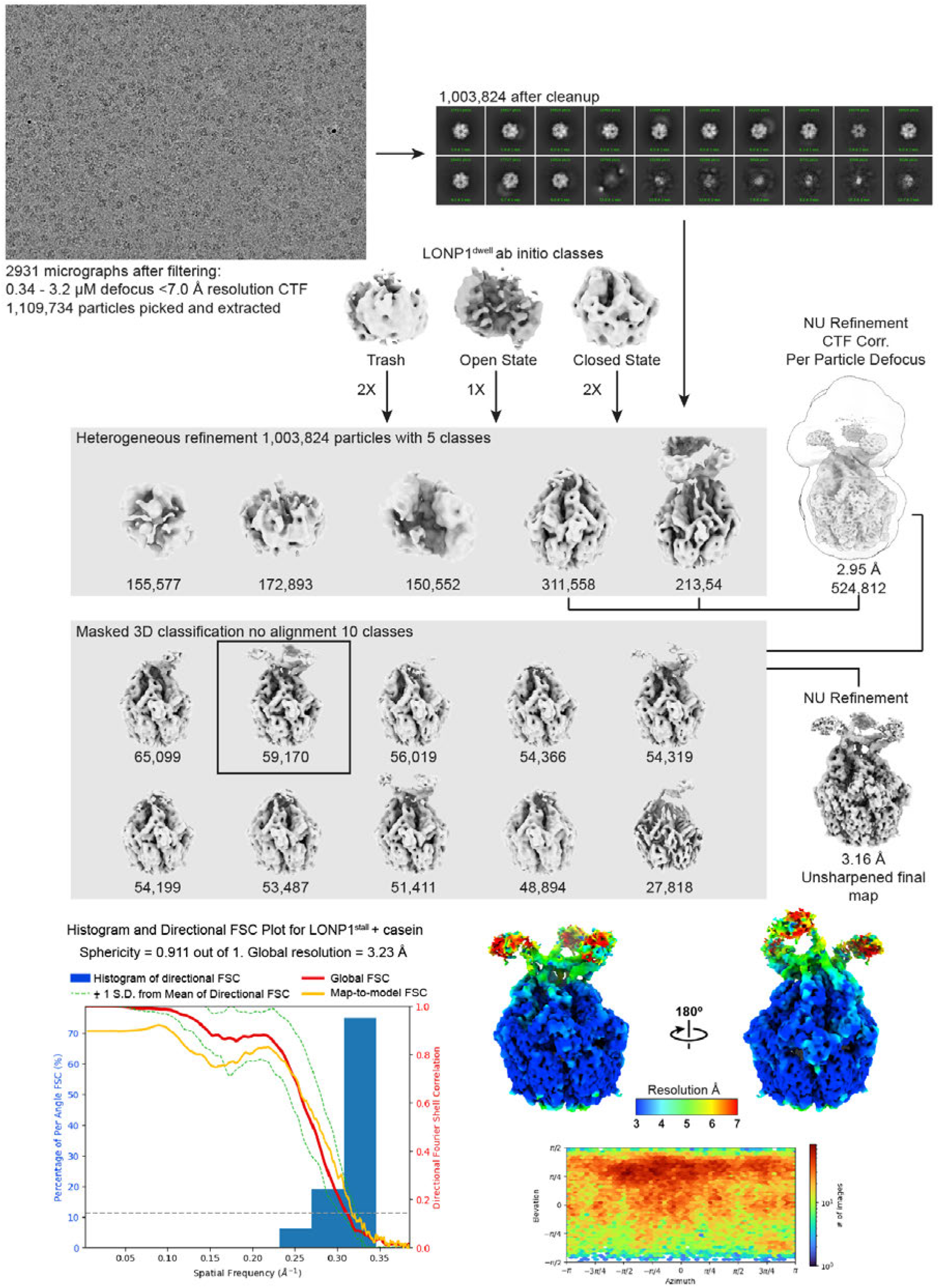
LONP1^stall^ + casein processing workflow.

**Figure S5.**
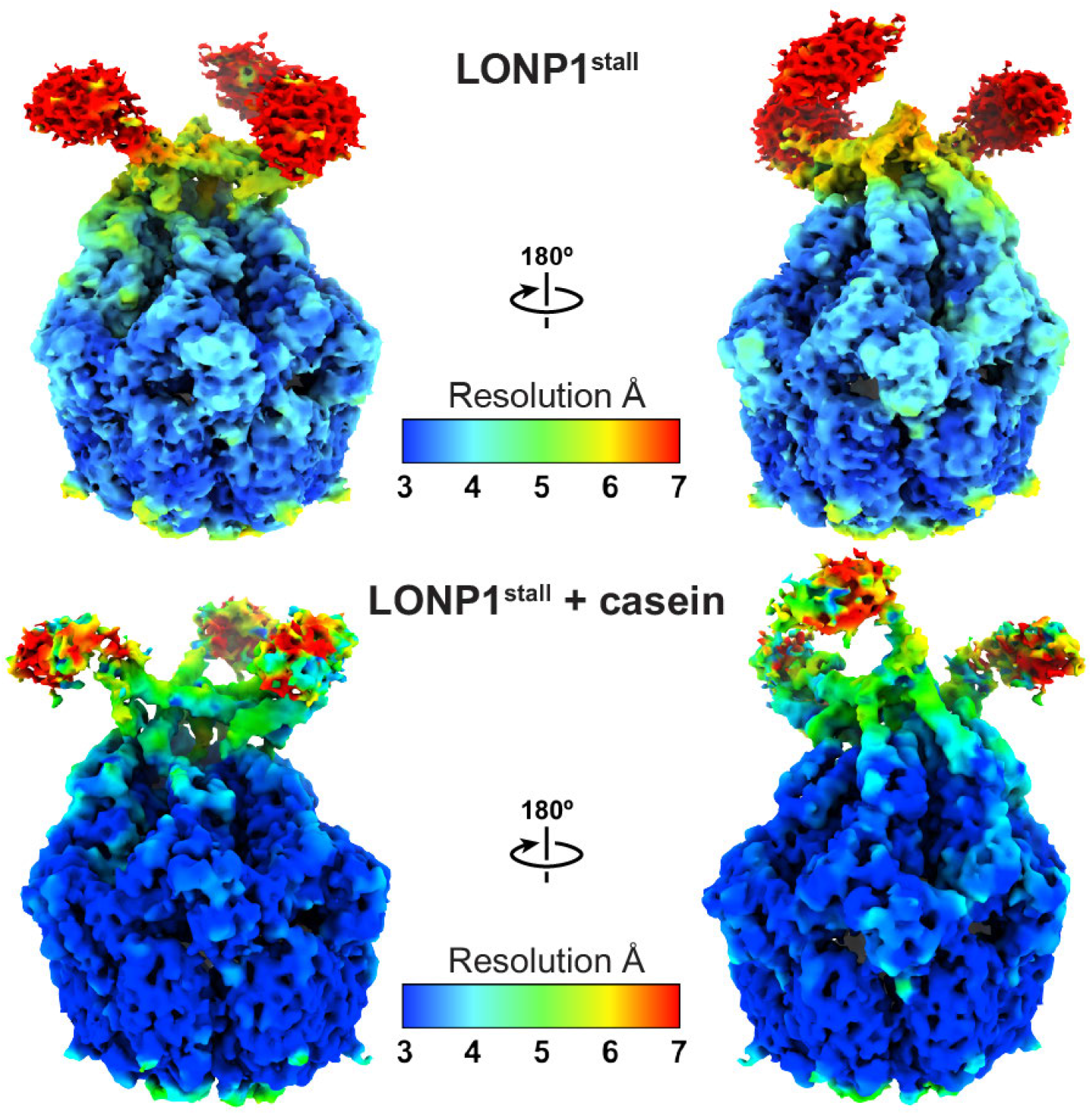
Comparison of local resolution of structures solved with and without the addition of casein. Binarization thresholds of LONP1^stall^ and LONP1^stall^ supplemented with casein are 0.0887 and 0.214, respectively.

**Figure S6.**
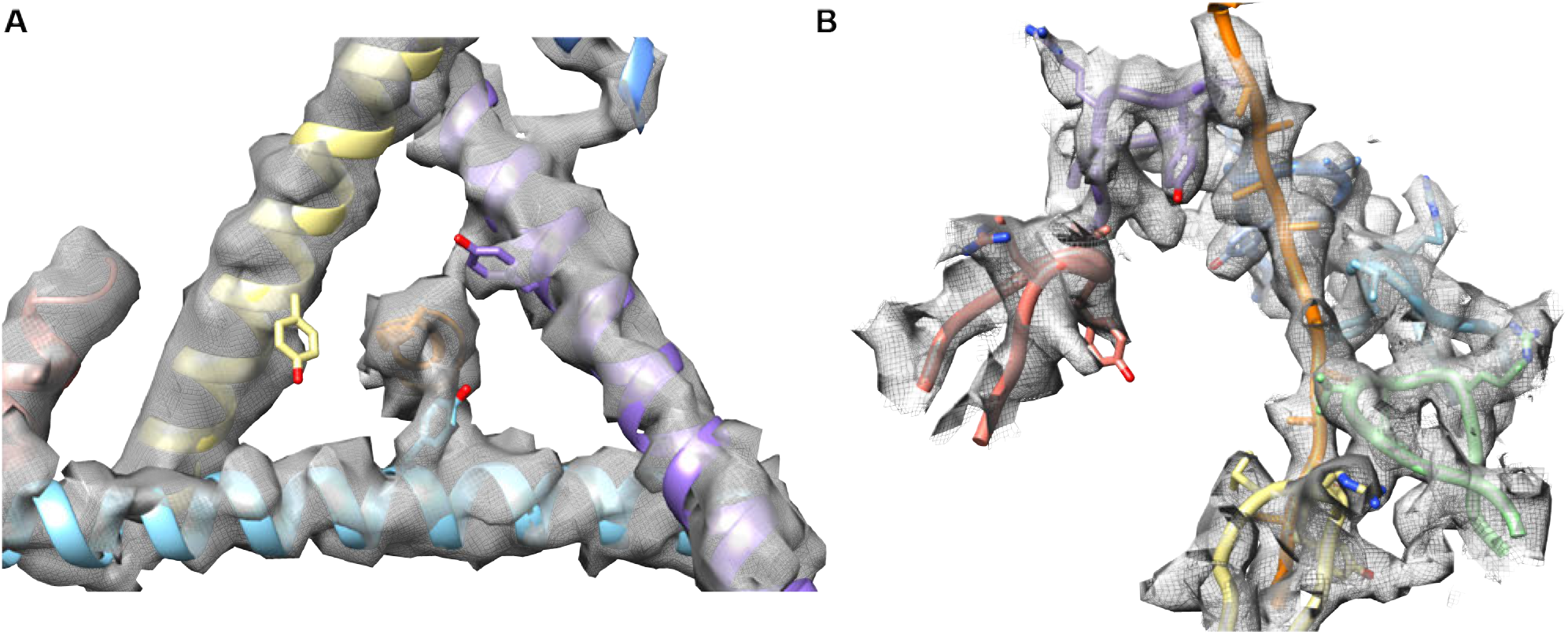
LONP1^stall^ CCD pore and pore loop EM density. **A**) CCD pore with Y394 shown as sticks interacting with bound substrate. Density shown from sharpened EM map with a 0.3 binarization threshold. **B**) Pore loop and substrate density. Density shown from sharpened EM map with a 0.375 binarization threshold.

**Figure S7.**
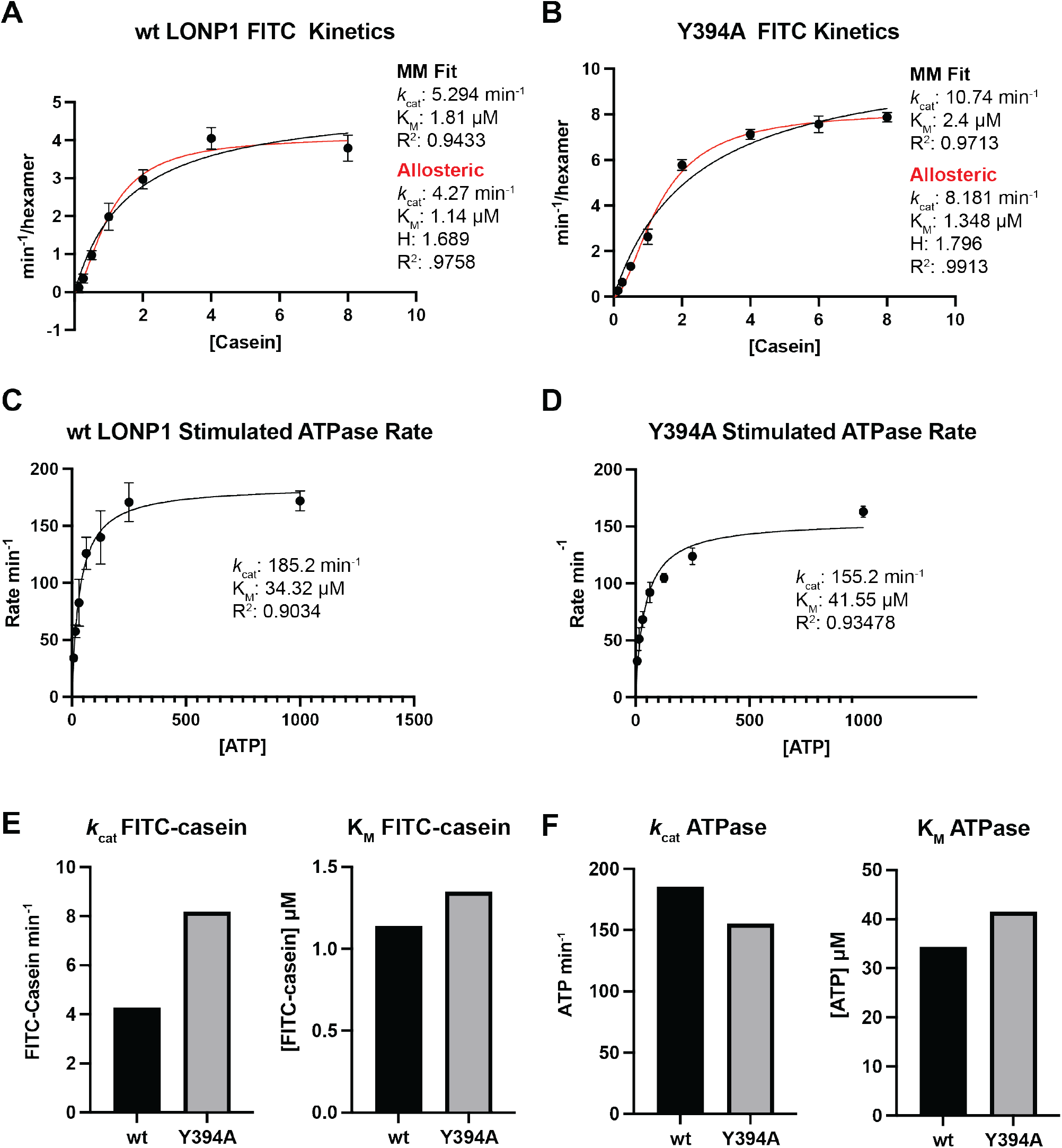
ATPase and FITC-casein degradation kinetics of wt LONP1 and Y394A mutant. **A**) FITC-casein degradation kinetics of wt LONP1. **B**) FITC-casein degradation kinetics of Y394A. **C**) Stimulated ATPase kinetics of wt LONP1. **D**) Stimulated ATPase kinetics of Y394A mutant. **E**) Comparative plot of the ATPase kinetic parameters (k_*cat*_ and K_*M*_) for wt LONP1 and Y394A mutant. **F**) Comparative plot of the FITC-casein degradation kinetic parameters (k_*cat*_ and K_*M*_) for wt LONP1 and Y394A mutant.

**Figure S8.**
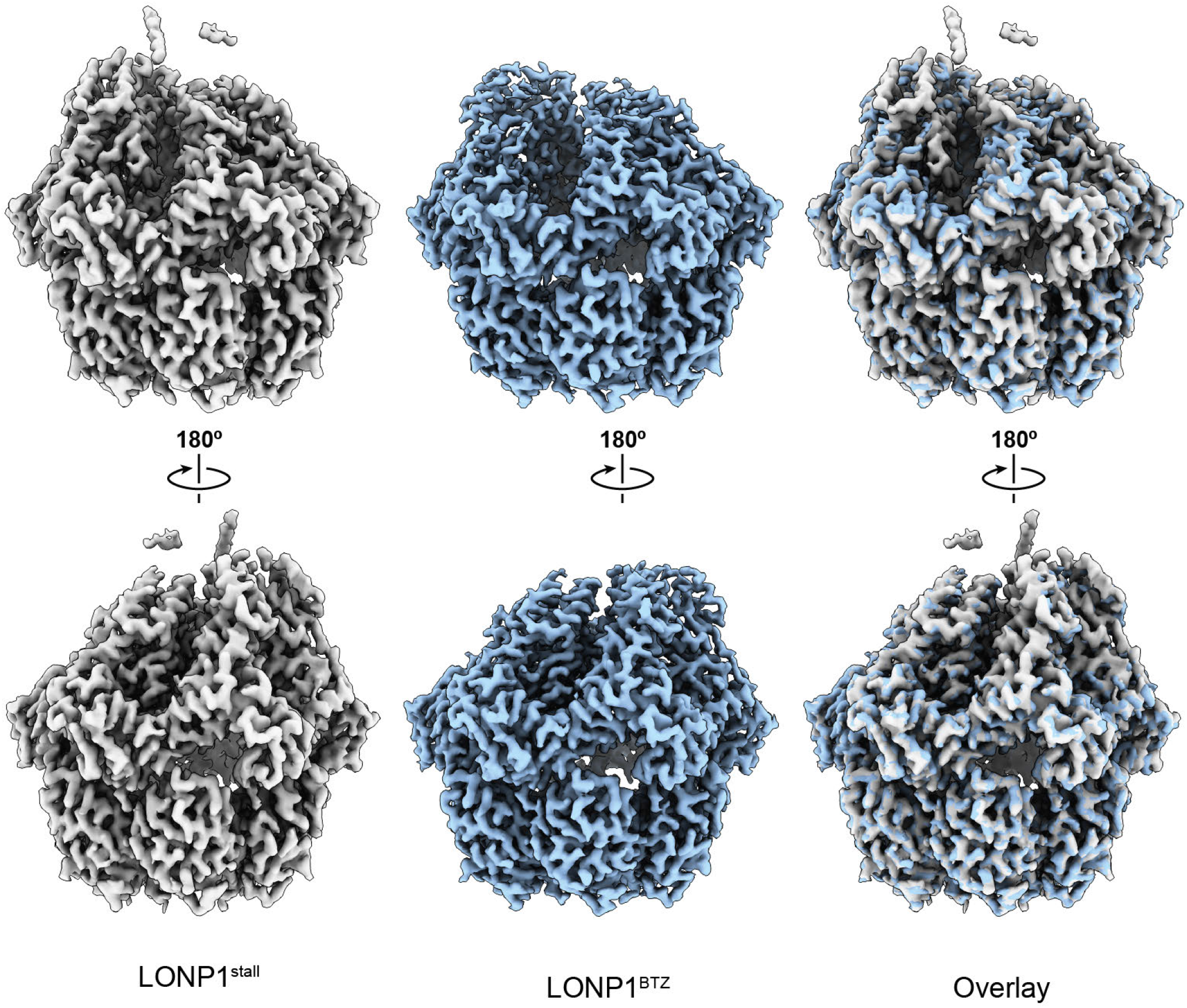
Comparison ADP-bound closed LONP1^stall^ and LONP1^BTZ^. Two viewing angles of LONP1^stall^, LONP1^BTZ^ (PDB ID: 7KRZ) and an overlay of LONP1^stall^ and LONP1^BTZ^ are shown from left to right, respectively. LONP1^stall^ and LONP1^BTZ^ adopt similar conformational states despite the lack of ATP in the LONP1^stall^ sample.

**Figure S9.**
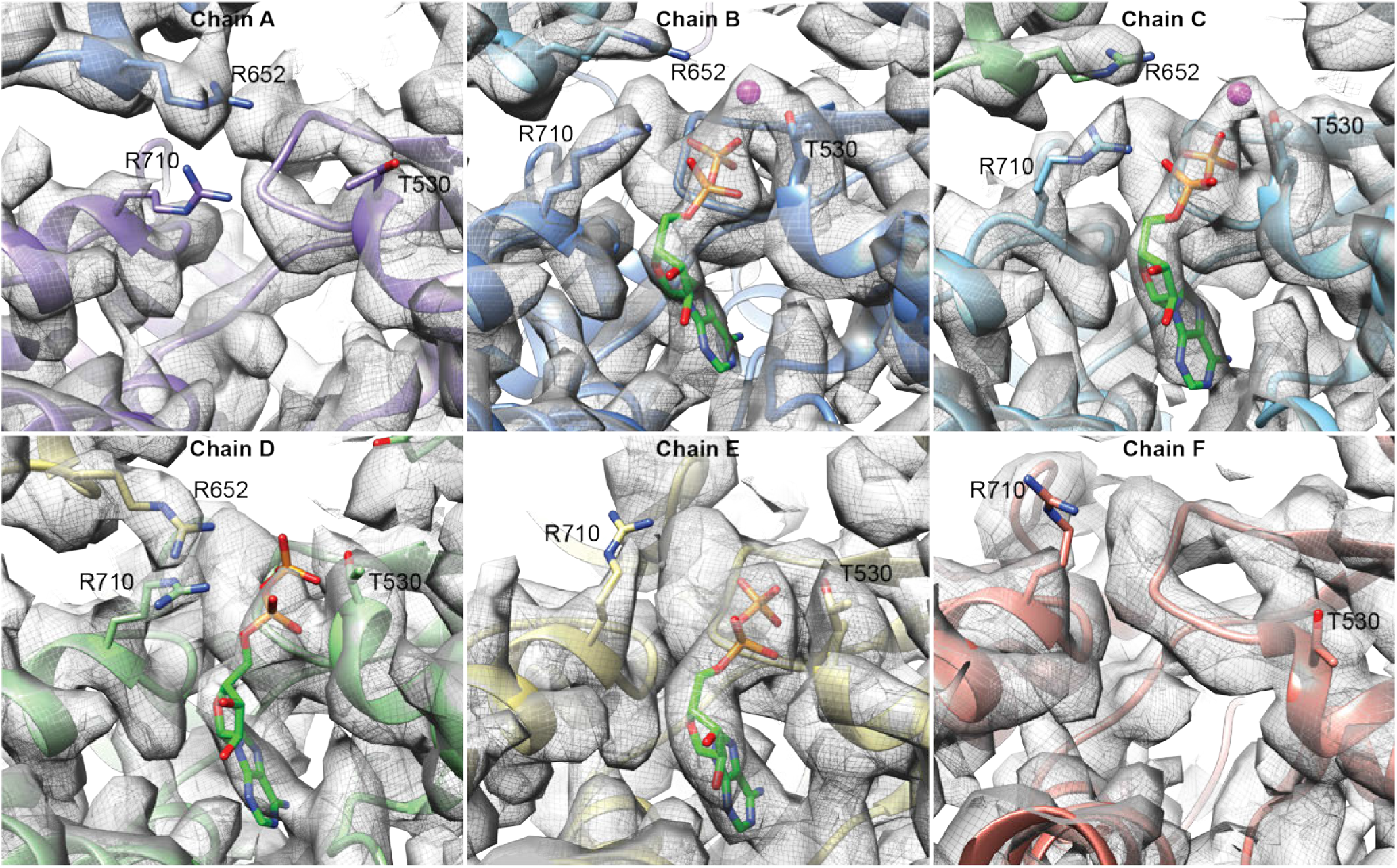
LONP1^stall^ nucleotide density. A binarization threshold of 0.4 was used for all panels. Chains A and F have no nucleotide density and only chains B and C had sufficient density to confidently model a magnesium ion.

**Figure S10.**
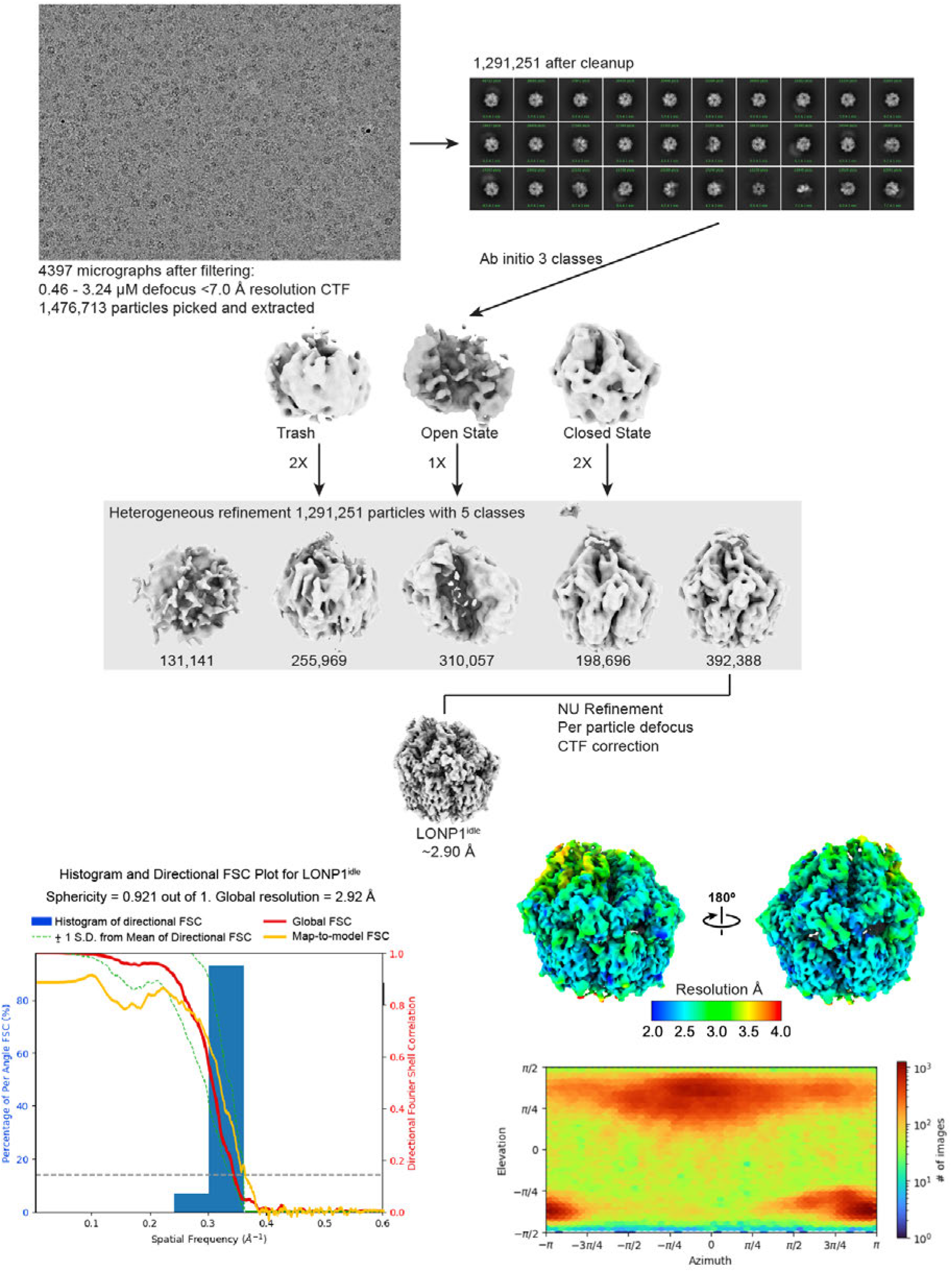
LONP1^idle^ processing workflow.

**Figure S11.**
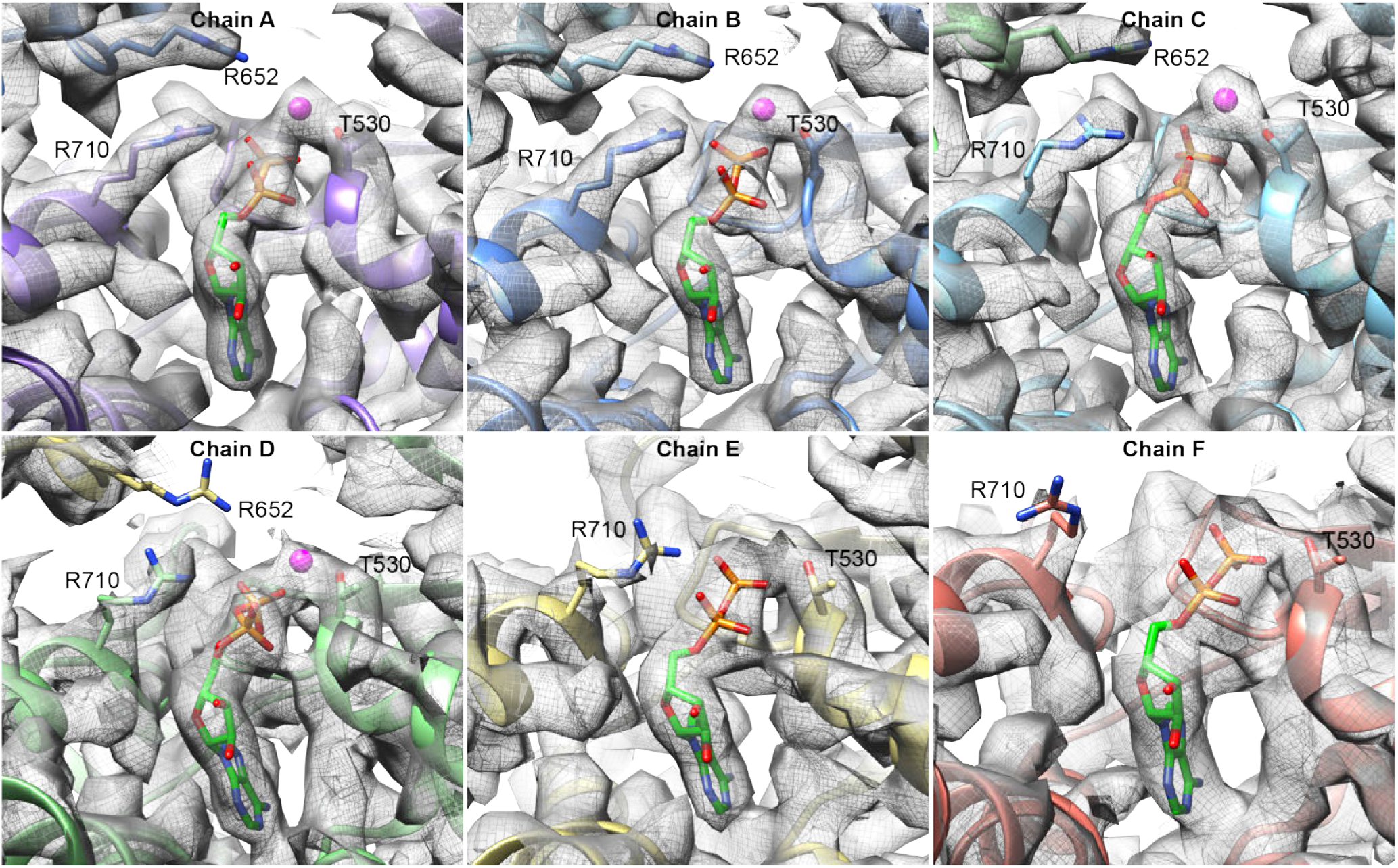
LONP1^idle^ nucleotide density. A binarization threshold of 0.6 was used for chains A-C and a threshold of 0.5 was used for chains E-F. Only chains A-D had sufficient density to confidently model a magnesium ion.

**Figure S12.**
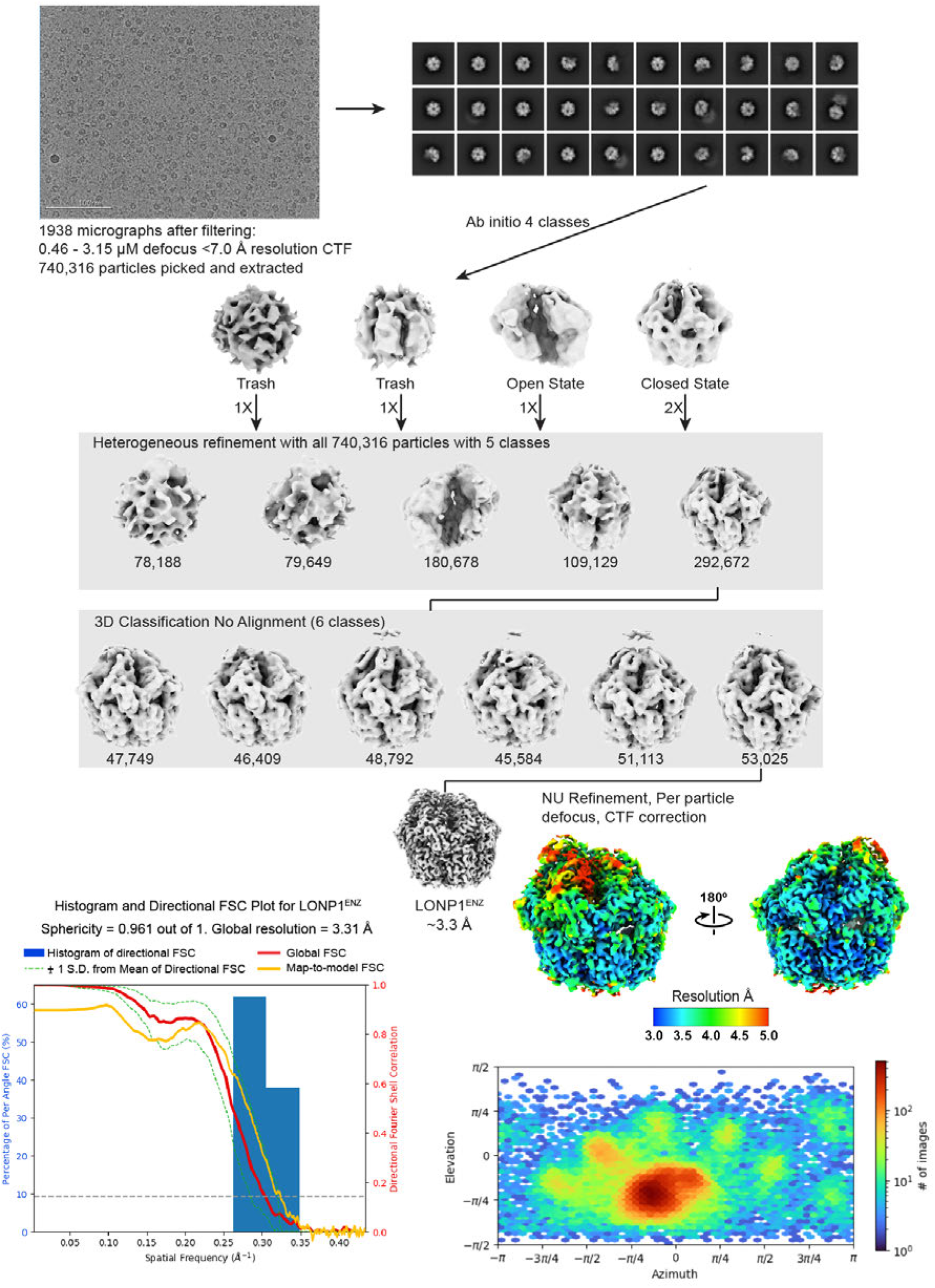
LONP1^ENZ^ processing workflow.

**Figure S13.**
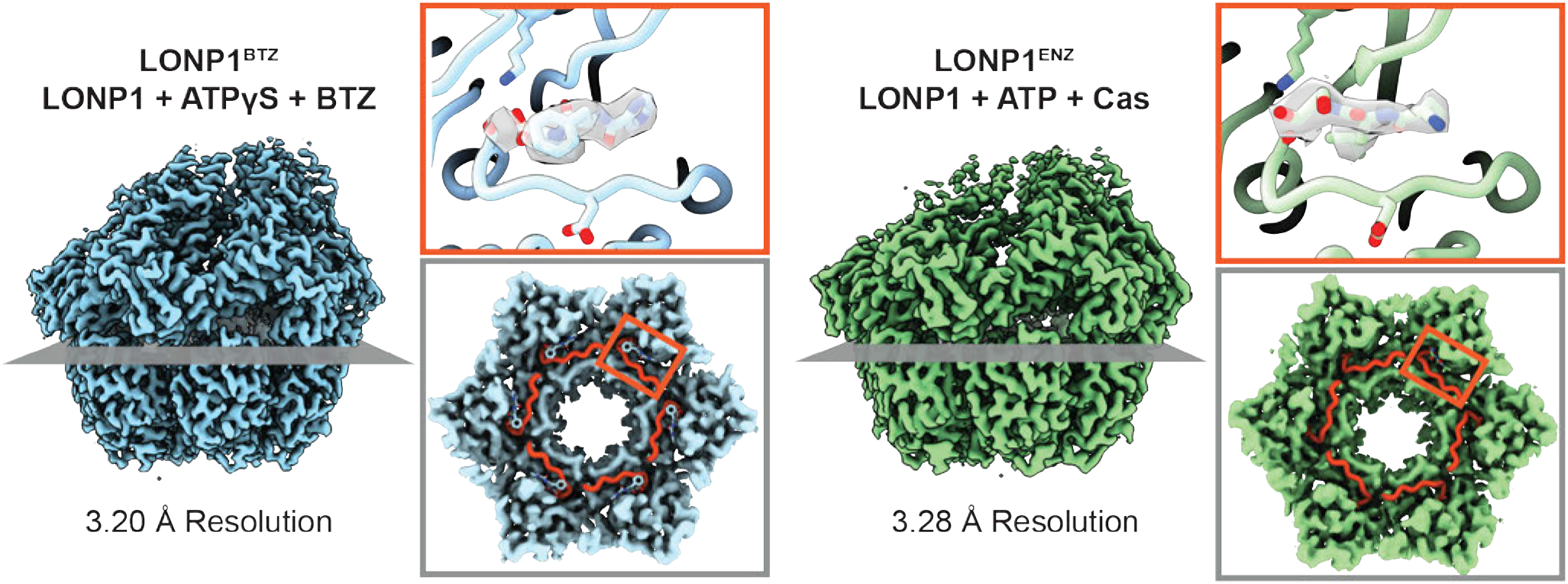
Comparison of LONP1^ENZ^ and LONP1^BTZ^. LONP1^BTZ^ (PDB ID: 7KRZ) is shown on the left and LONP1^ENZ^ shown on the right. The gray cross section in both structures corresponds to the grey outlined panel where the inhibitory loop (colored red) is in the open active conformation for every active site. The red box corresponds to the red outlined panel demonstrating the EM density for either bortezomib or peptide bound in the proteolytic active site.

**Figure S14.**
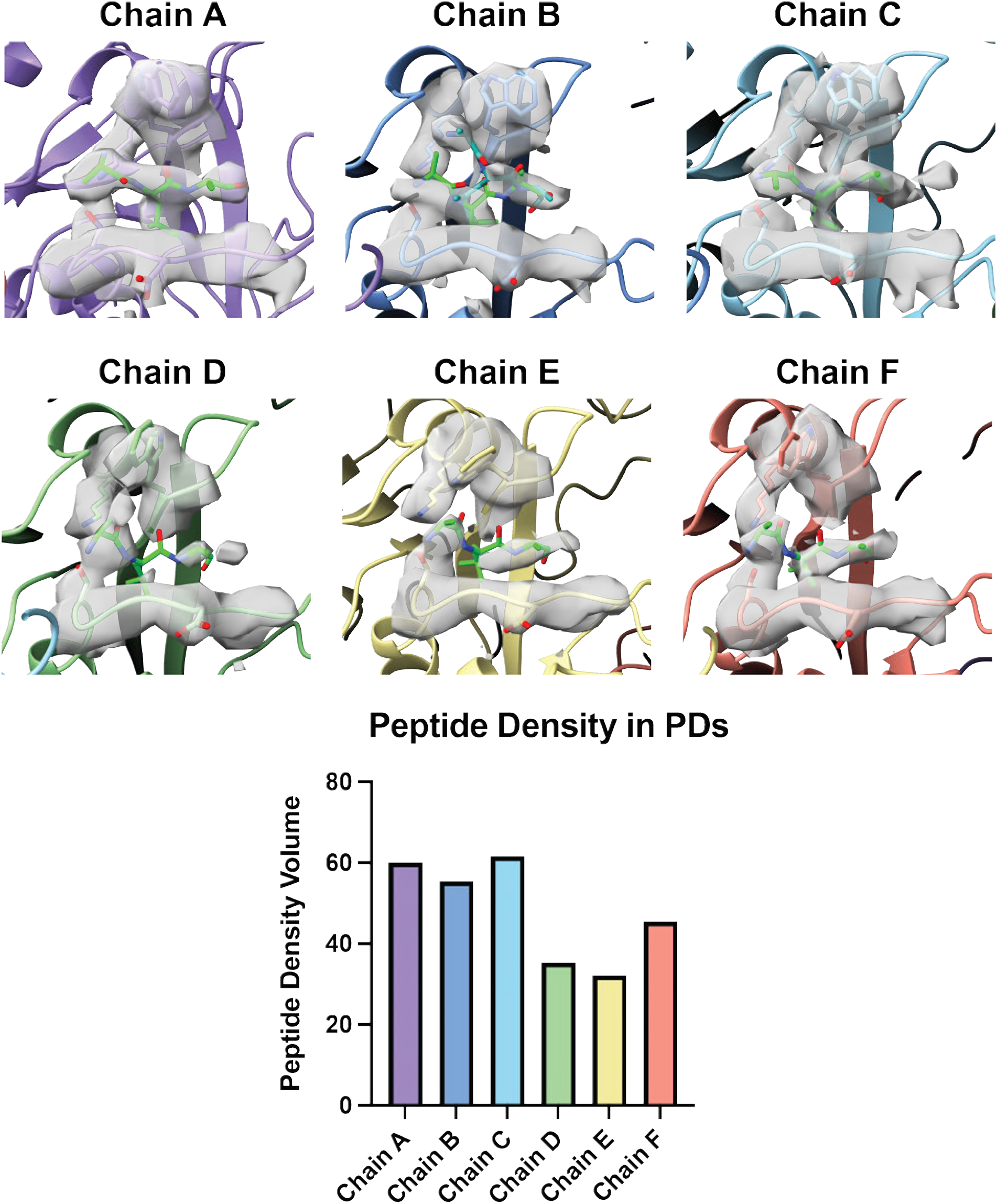
Protease domain substrate density analysis. A tripeptide (Ala-Leu-Ala) was modelled into each protease domain active. The peptide density decreases as a function of position in the helical register. Positions A-C have more defined substrate density with density that accommodates the central leucine residue of the tripeptide. However, positions D-F have notably weaker signal for bound substrate. The area associated with bound substrate was calculated with the same method used to determine γ-phosphate density (see *Methods*).

**Figure S15.**
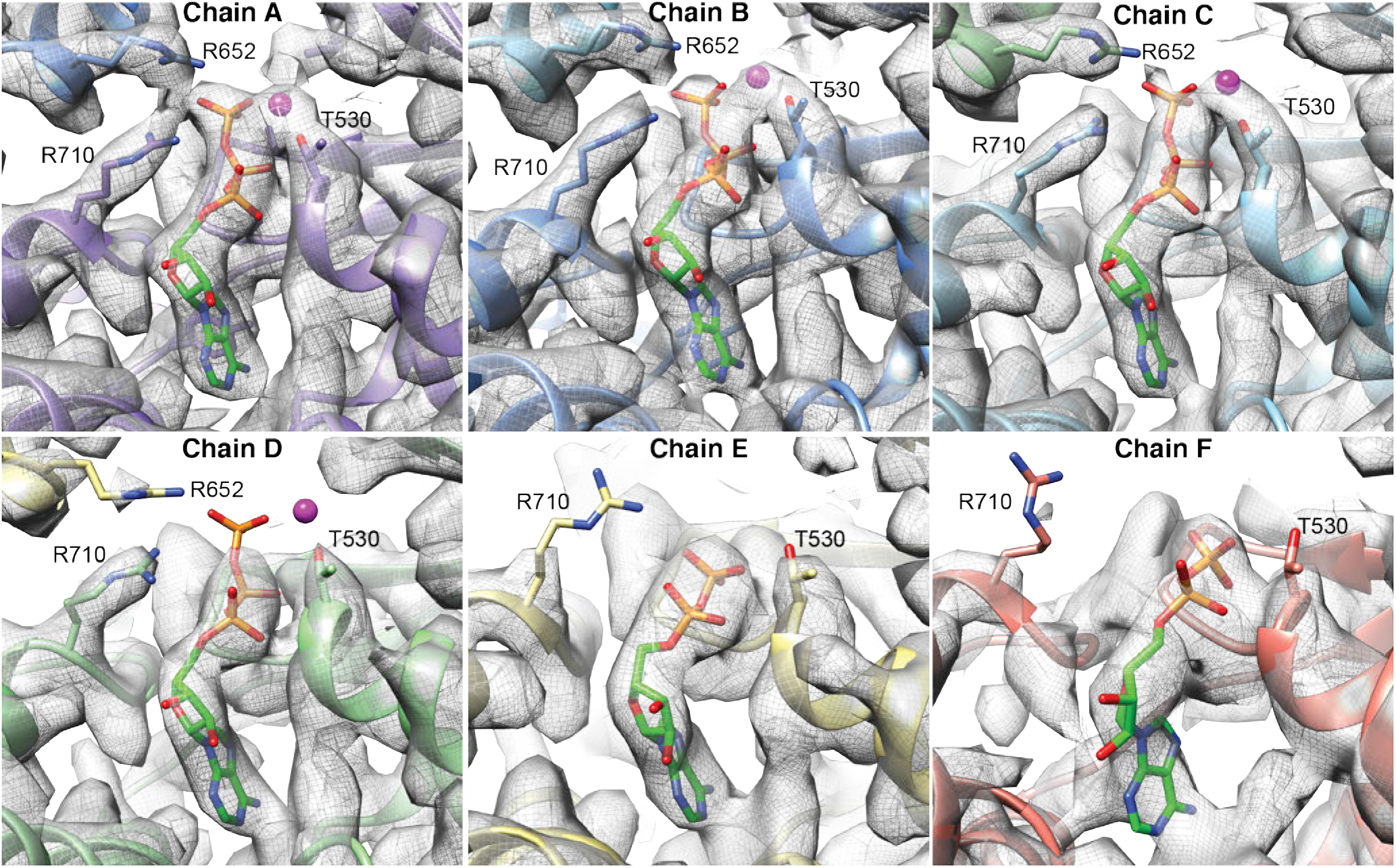
LONP1^ENZ^ nucleotide density. Map was contoured at a binarization threshold of 0.35 for all panels.

**Table S1.** Cryo-EM data collection, analysis, and modeling statistics.

|  | LONP1<sup>stall+cas</sup> | LONP1<sup>idle</sup> | LONP1<sup>ENZ</sup> | LONP1<sup>stall</sup> |
| --- | --- | --- | --- | --- |
| EMDB ID | EMD-45433 | EMD-45431 | EMD-45430 | EMD-45434 |
| PDB ID | 9CC3 | 9CC1 | 9CC0 |  |
| **Data Collection** |  |  |  |  |
| Microscope | Titan Krios | Titan Krios | Talos Arctica | Talos Arctica |
| Voltage (kV) | 300 | 300 | 200 | 200 |
| Camera | K3 BioQuantum | K3 BioQuantum | K2 Summit | K2 Summit |
| Camera mode | Counting | Counting | Counting | Counting |
| Magnification (nominal/at detector) | 105,000/60,024 | 105,000/60,024 | 36,000/43,478 | 36,000/43,478 |
| Data acquisition software | Leginon | Leginon | Leginon | Leginon |
| Exposure navigation | Image shift to 49 holes | Image shift to 49 holes | Image shift to 16 holes | Image shift to 16 holes |
| Total electron exposure (e<sup>-</sup>/Å<sup>2</sup>) | 54 | 54 | 50 | 50 |
| Exposure rate (e<sup>-</sup>/pixel/sec) | 30 | 30 | 6.75 | 6.87 |
| Frame length (ms) | 20 | 20 | 200 | 200 |
| Number of frames | 50 | 50 | 49 | 48 |
| Pixel size (Å/pixel)* | 0.833 | 0.833 | 1.15 | 1.15 |
| Defocus range at scope (μm) | 1 | 1 | 0.9 - 1.16 | 0.9 - 1.6 |
| Measured defocus range (μm) | 0.32 - 3.28 | 0.46-3.24 | 0.46-3.15 | 0.10 - 2.48 |
| Energy filter slit width (eV) | 20 | 20 | N/A | N/A |
| Micrographs collected (no.) | 2,931 | 4,397 | 1,938 | 1,944 |
| **Data analysis** |  |  |  |  |
| Image processing package | cryoSPARC | cryoSPARC | cryoSPARC | cryoSPARC |
| Total extracted picks (no.) | 1.11M | 1.47M | 740,316 | 496,793 |
| Particles used for 3D (no.) | 524,812 | 591,084 | 292,672 | 229,101 |
| Final Particles (no.) | 59,170 | 392,388 | 53,025 | 94,000 |
| Symmetry | C1 | C1 | C1 | C1 |
| Resolution (global) (Å) |  |  |  |  |
| FSC 0.5 (unmasked / masked) | 7.23 / 3.82 | 3.87 / 3.24 | 7.77 / 3.81 | 6.14 / 3.55 |
| FSC 0.143 (unmasked / masked) | 3.89 / 3.28 | 3.31 / 2.89 | 4.0 / 3.28 | 3.65 / 3.17 |
| Local resolution range (Å) | 2.9 – 38.2 | 1.8 – 24.6 | 2.8 – 47.3 | 2.7 – 48.9 |
| 3DFSC Sphericity (%) | 91.0 | 92.0 | 96.1 | 98.1 |
| Map sharpening *B*-factor (Å<sup>2</sup>) | -67.5 | -90 | -60 | -60 |
| **Model composition** |  |  |  |  |
| Protein residues | 3353 | 3135 | 3140 |  |
| Ligands | 8 | 12 | 10 |  |
| **Model Refinement\*** |  |  |  |  |
| Refinement package | ISOLDE/Phenix | ISOLDE/Phenix | ISOLDE/Phenix |  |
| MapCC (volume/mask) | 0.81 / 0.82 | 0.83 / 0.84 | 0.85/ 0.85 |  |
| B-factors (Å<sup>2</sup>) |  |  |  |  |
| Protein residues | 80.19 | 47.63 | 86.73 |  |
| Ligands | 75.67 | 44.93 | 82.86 |  |
| R.m.s. deviations |  |  |  |  |
| Bond lengths (Å) | 0.002 | 0.003 | 0.003 |  |
| Bond angles (°) | 0.528 | 0.589 | 0.535 |  |
| **Validation** |  |  |  |  |
| Map-to-model, FSC 0.5 | 3.42 | 3.08 | 3.40 |  |
| Ramachandran plot |  |  |  |  |
| Favored (%) | 98.61 | 98.29 | 98.16 |  |
| Allowed (%) | 1.39 | 1.71 | 1.84 |  |
| Outliers (%) | 0.0 | 0.0 | 0.00 |  |
| MolProbity score | 1.23 | 1.29 | 1.84 |  |
| Clashscore | 4.57 | 5.36 | 5.40 |  |
| Poor rotamers (%) | 0.00 | 0.04 | 0.15 |  |
| C-beta deviations | 0 | 0 | 0 |  |
| CaBLAM outliers (%) | 1.00 | 0.75 | 0.88 |  |
| EMRinger score | 2.26 | 3.06 | 2.83 |  |
| Qscore | 0.489 | 0.538 | 0.501 |  |

